# Fixation shifts in a novel “no-report” binocular rivalry paradigm induce saccade-related perceptual switches

**DOI:** 10.1101/2025.02.09.637085

**Authors:** Ege Kingir, Ryo Segawa, Janis K. Hesse, Igor Kagan, Melanie Wilke

## Abstract

No-report paradigms help to avoid report-related confounds in conscious perception studies. A novel no-report binocular rivalry paradigm by Hesse and Tsao (2020) tracks conscious content using eye position as subjects follow fixation points linked to the rivaling stimuli. However, it remains unclear whether perceptual changes arise spontaneously or are induced by external factors such as fixation shifts and saccades. We found an increased probability of perceptual switches time-locked to fixation point shifts, indicating that some switches are externally driven. To disentangle the effects of visual fixation shifts and saccades, we implemented a two-factorial design and found that saccades play a larger role in eliciting perceptual changes. We estimate that 14% of saccades trigger a switch, accounting for 24% of all perceptual transitions. Our findings provide an analysis framework and guidelines for excluding externally driven perceptual switches, enabling a clearer focus on internally generated perceptual dynamics.

## 1. INTRODUCTION

Which neural mechanisms underlie our phenomenal, conscious experience is one of the most fascinating and unresolved questions in neuroscience (Bachmann & Aru, 2023; Lepauvre & Melloni, 2021; Storm et al., 2024). Identifying the neural correlates of consciousness (NCC) is not only important from a philosophical and basic scientific point of view, but has also clinical implications, as it would enable judgments about the contents of awareness in severely brain-damaged patients (Dehaene & Changeux, 2011; Storm et al., 2017). Subjective perception is often probed with optical illusions where a constant physical stimulus elicits different percepts. A popular approach to study fluctuating visual perception with a constant physical input is the use of binocular rivalry paradigms. Binocular rivalry emerges when two dissimilar images (e.g., house or face) are presented monocularly to different eyes. Instead of creating a fusion of the images, the brain instantiates a perceptual competition between the two stimuli such that typically only one of the stimuli is perceived at a given time, with piecemeal perception during transition phases, despite constant visual input (Carmel et al., 2010; Carter et al., 2020; Tong et al., 2006).

The neural correlates of those subjective visual states are typically referred to as ‘neural correlates of conscious perception’ as the physical stimulus remains constant. Binocular rivalry has been studied in a wide range of brain areas and at different spatial and temporal scales, employing diverse methods such as single cell and local field potential (LFP) recordings, electroencephalographic recordings (EEG / MEG) and functional MRI in monkeys and humans (Storm et al., 2017; Tong et al., 2006; Zhang et al., 2017). Some of those studies either focused on the perceptual dominance versus suppression phases of a given stimulus, while others focused on the neural underpinnings of the perceptual switch process itself (Dwarakanath et al., 2023). As a general pattern, the electrophysiological and fMRI studies focusing on stimulus perception (‘content’) found a gradual increase of dominance/suppression-related activity from early stages of visual processing to higher-order object-selective regions (i.e. LGN / V1 < extrastriate visual cortex (V4/MT) < inferotemporal cortex) (Gail et al., 2004; Gelbard-Sagiv et al., 2018; Hesse & Tsao, 2020; Keliris et al., 2010; Leopold & Logothetis, 1996; Logothetis & Schall, 1989; Maier et al., 2007; Panagiotaropoulos et al., 2012; Sheinberg & Logothetis, 1997; Tong et al., 2006). Studies which focused on the perceptual switch process typically reported stronger activity in parieto-frontal regions when (monocular) physical stimulus switches were compared to internally induced ones (Brascamp et al., 2018; Dwarakanath et al., 2023; Kornmeier & Bach, 2012; Leopold & Logothetis, 1999; Lumer et al., 1998; Pettigrew, 2001).

However, one major problem remained with the aforementioned ‘neural correlates of consciousness’ on both theoretical and empirical grounds. Since bistable perception is subjective, experimenters need to rely on explicit perceptual reports from the subjects such as button responses to track perceptual switches. These reports confound the NCC with many other cognitive processes such as resolving visual complexity, attentional factors, working memory, motor planning and performance monitoring (Aru et al., 2012; Tsuchiya et al., 2015). Indeed, when the behavioral report was removed, and perception of a bistable stimulus was inferred either from subtle stimulus manipulations or from physiological measures such as small eye movements and pupil diameter, perceptual modulation in certain frequency bands disappeared and switch-related activity in fronto-parietal regions was diminished (Andersen et al., 2022; Frässle et al., 2014; Kloosterman et al., 2015; Wilke et al., 2009).

Thus, to eliminate report-related confounds, many consciousness researchers advocated for adding so called ‘no-report paradigms’ to the study of visual consciousness (Duman et al., 2022; Tsuchiya et al., 2015, 2016). For binocular rivalry, several clever no-report approaches have been developed, based on electrophysiological frequency tagging of the stimuli, stimulus contrast manipulations in combination with pupil size assessments and optokinetic nystagmus (reviewed in: Duman et al., 2022). In each case, the validity of the no-report perceptual readout was established before by comparing the results inferred from the no-report condition with those obtained from explicit reports in humans. For instance, previous studies using optokinetic nystagmus presented the subjects (monkeys or humans) with drifting gratings moving in opposite directions in each eye. They first established in monkeys and humans that the slow phase of the optokinetic nystagmus correlates tightly with conscious reports of perceptual dominance during binocular rivalry as assessed by saccade or lever reports (Frässle et al., 2014; Logothetis & Schall, 1989; Naber et al., 2011).

For example, in the Frässle et al. fMRI study (2014), the timing of perceptual switches was inferred from changes in the direction of nystagmus in the presence and absence of explicit reports from subjects. Even in the absence of reports, subjects continued to experience vivid rivalry. In both report and no-report condition, occipital and parietal areas showed differential activity in correlation with the inferred perceptual contents. However, the same correlation that was present in the report condition in frontal areas disappeared in the absence of report. Similar results were obtained using pupil size as a perceptual readout during binocular rivalry between static gratings of low and high contrast (Frässle et al., 2014). These studies point to the possibility that previously reported switch-related activity in the medial frontal gyrus is caused by factors such introspection and self-monitoring associated with the difficulty of reporting ambiguous percepts, but not directly related to switching between conscious percepts *per se*. It should be noted however, that recent single cell and local field potentials studies in dorsolateral prefrontal cortex in rhesus macaques that used OKN as perceptual no-report readout, found not only feature-selective perceptual modulation and robust single trial decoding of perceptual contents (Kapoor et al., 2022; Panagiotaropoulos et al., 2012) but also state fluctuations in low frequency bands predicting perceptual reversals (Dwarakanath et al., 2023).

On the other hand, OKN might not be a perfect no-report paradigm for all NCC-related questions. For example, the use of drifting stimuli is not appropriate for the use with natural images or not when eye movement direction presents a major confound in a given brain region. Moreover, on the retina, the stimulus in one eye will be mostly stationary, as it is being followed by the OKN, and the stimulus in the other eye will be drifting with double velocity in the opposite direction of the OKN, causing a physical difference between the two percept conditions. Another questionable aspect of OKN as a perceptual readout is the fact that spontaneous OKN fluctuations during binocular rivalry can still be obtained in ketamine-anaesthetized monkeys, casting doubt on the perfect relation between OKN and the contents of conscious perception (Leopold et al., 2002). Other no-report techniques such as pupil size responses are also not perfect as pupil size is affected by many other factors such as luminosity, general arousal, surprise and confidence (Joshi & Gold, 2020; Kloosterman et al., 2015; Lempert et al., 2015; Urai et al., 2017).

A new promising no-report binocular rivalry method was recently developed by Hesse and Tsao to decode the conscious percept from face patches in the inferotemporal cortex of macaques (Hesse & Tsao, 2020). The advantage of this paradigm is the flexible use of non-moving binocular rivalry stimuli, while the subjects have an orthogonal task of following a moving fixation spot to not get bored. The paradigm uses two stimuli (e.g., a face and an object), both of which incorporate their own fixation point (in the same color as the corresponding stimulus), which changes position every 2000 ms in humans and 800 ms in the monkeys. When perceiving the face, subjects will generally perceive the fixation spot in the corresponding eye of the face, and saccade to it, and vice versa when perceiving the object. The consciously perceived object at each time point can thus be inferred from the distance of the gaze location to each fixation point. Hesse and Tsao showed that large proportions of IT neurons represented the conscious percept even without active report. Furthermore, on single trials, they could decode both the conscious percept and the suppressed stimulus (Hesse & Tsao, 2020).

The novel fixation paradigm arguably fixes some problems of the OKN approach, such as restrictions in the type of stimuli that can be used, and is well suited to investigate the neural correlates of perceptual states with constant physical stimuli. The aim of our current study was to test the suitability of this paradigm for the investigation of neural correlates of the perceptual switch process itself. Specifically, we tested whether the sudden fixation point shifts and/or the ensuing eye movement can trigger a perceptual switch in this no-report paradigm. In the original BR paradigm from Hesse and Tsao (2020) subjects need to follow the fixation point they are perceiving, and once their perception switches from one stimulus to the other, subjects will find the fixation point of this new object and saccade to this new gaze location. Both fixation points change location regularly to keep the subjects engaged in the task and actively track the fixation point of the perceived object. In line with a previous finding from Van Dam and van Ee (2006) that saccades facilitate perceptual switches, we hypothesize that the process of planning and performing a saccade to the jumping fixation points may induce perceptual switches that the subject otherwise would not spontaneously experience. Furthermore, abrupt appearances of additional visual probes on top of the rivaling stimuli modulate perceptual dominance durations to favor longer-lasting perception of the image on which they are shown (Metzger & Beck, 2020). In Hesse and Tsao’s paradigm, fixation points corresponding to both images change location every 2 seconds, resembling an onset of new visual probes on rivaling stimuli. We hypothesize that this external factor of stimulus novelty may further induce perceptual switches that would not occur if the stimuli were kept stable.

To test these hypotheses, we first analyze perceptual changes time-locked to the onset of fixation point shifts (and naturally also time-locked to the following saccade) in the original variant of the Hesse and Tsao paradigm. In a second step, we add three additional tasks to dissociate the effects of fixation point shifts and saccades on induced perceptual switches. We show that saccades play a larger role in facilitating perceptual switches than the visual transients due to fixation point shifts. We also discuss possible adaptations of this promising no-report paradigm, and propose analysis frameworks for the investigation of spontaneous perceptual switch processes.

## 2. METHODS

### 2.1. Participants

Twenty healthy subjects (6 male, 14 female, mean age: 26.3, age range: 22-34) participated in this experiment. Subjects had normal or corrected-to-normal vision, normal hearing, unrestricted arm- and hand-mobility, no neurological or psychiatric disorders, no recent use of drugs, medications or alcohol dependence. At the beginning of the first session, subjects signed an informed consent form, which included basic information about the tasks that they are required to perform. They were compensated with 15 Euro/hour for their participation. Each subject visited the laboratory on two different days in order to complete all four tasks, and the order of tasks for each subject was pseudorandomized. Three subjects had to be excluded from data analysis, Subject #03 for performing much lower number of trials than any other subject in one of the tasks (see **Supplementary Table 1**), and Subject #05 and #07 for not being able to attend their second experiment sessions.

Experiments were performed in accordance with institutional guidelines for experiments with humans and adhered to the principles of the Declaration of Helsinki. The experimental protocol was approved by the ethics committee of the Georg-Elias-Mueller-Institute for Psychology, University of Goettingen (GEMI 17-06-06 171).

### 2.2. Experimental Setup and Stimulus Properties

Testing procedures took place in a dark room at the Germany Primate Center (DPZ). Subjects sat 47 cm away from the center of the screen, while resting their head on a chin and forehead rest, and performed the tasks while wearing cyan-red 3D anaglyph glasses. Stimuli were presented on an LCD monitor with a refresh rate of 120 Hz and 2560×1444 pixels resolution (M27Q Gaming Monitor (GIGABYTE, Taipei City, Taiwan)). Rivaling stimuli size was 7.5 visual degrees (°) in the vertical and 5.8° in the horizontal dimension, centered on the middle of the screen. Each fixation point was 0.3°, and the ‘peripheral’ fixation points corresponding to each of the rivaling stimuli (used in tasks 1 and 3, see **Fig. 1B**) were 1.5 visual degrees away from the center of the screen. Visual stimuli were generated with MATLAB (The MathWorks, Inc, Natick, MA) and the Psychophysics Toolbox (Brainard, 1997), using the custom toolbox monkeypsych (https://github.com/dagdpz/monkeypsych).

**Figure 1:**
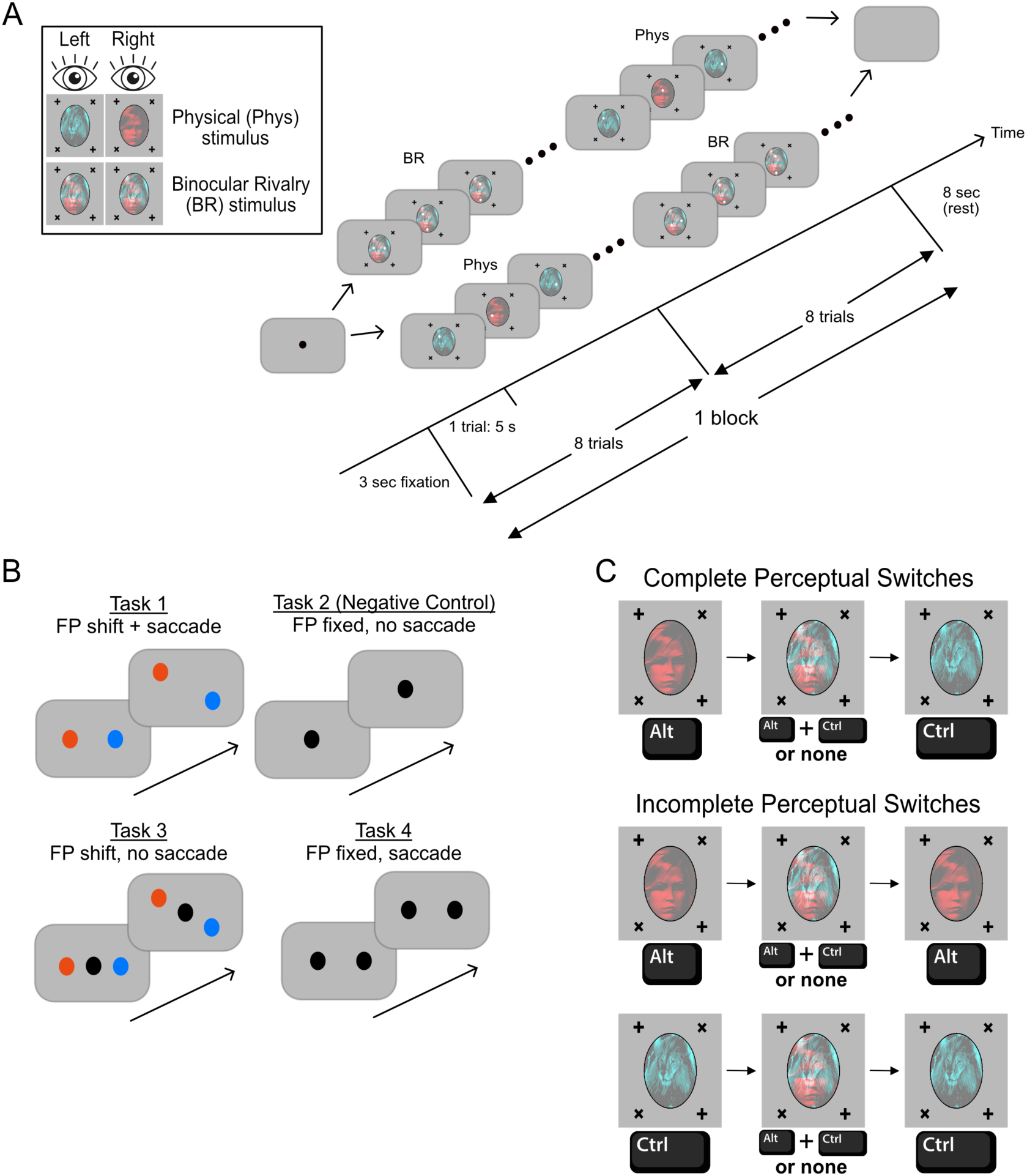
Experiments and Task Designs: **(A)** Stimulus identities and block structure, common to all tasks: Binocular Rivalry trials (‘BR’) and Physical switch trials with monocularly presented targets (‘PHYS’) conditions (8 trials each) are presented alternatingly in a pseudorandomized order, constituting a block of 16 trials, which in turn is repeated 16 times for each task. A trial is 5 seconds for all tasks. **(B)** Differences between the four tasks: In Task 1, subjects were instructed to fixate on the fixation point (FP) they see. The location of each FP changed every 5 seconds, requiring the subject to find and fixate on the new location of the FP. In Task 2, fixation point remained stable and 8 trials were continuous. In Task 3, FPs changed location every 5 seconds but subjects were instructed to keep fixating to the central FP (black). In Task 4, an auditory cue told subjects to change their fixation from one black FP to the other. **(C)** Possible types of perceptual switches and the button(s) that subjects pressed to report each percept. Perceptual switches in each task were sometimes completed, such that perceived stimulus switched from one to the other smoothly. However, there were also some cases where the perceived stimulus went back to the old one, following a piecemeal perception period. We were able to distinguish between the two types of switches according to the order of the button release-press pattern associated with each switch type. The stimulus material is from one of the authors (Hesse, J. K., & Tsao, D. Y. (2020), *eLife*).

Eye movements were tracked with a monocular head-mounted miniature ViewPoint eye tracker (Arrington Research, Inc., Scottsdale) placed below the eye and behind the anaglyph glasses. Gaze position was sampled continuously with a temporal resolution of 60 Hz. Before the start of the experiment, the eye tracker was calibrated using a 4 by 5 points calibration matrix using the ViewPoint piece-wise linear calibration, and converted from raw camera view-based gaze data to degrees of visual angle using linear transformation with independent gains and offsets for the vertical and horizontal positions.

Before each session, we performed a stimulus calibration session to balance rivaling stimuli perceived contrasts in overlaid cyan and red objects, in which the subject reported their perceived color (CTRL button on the keyboard for perceived cyan object - lion, and ALT button for the red object - face), while fixating on a spot in the middle of the screen. Subjects performed these continuous calibration sessions (up to 5 minutes) until the two stimuli were perceived for similar durations (within 40% to 60%). If the subject reported that the fixation point of either the red or the cyan stimulus is “always visible” or “always invisible” (irrespective of the perceived object), we adjusted the fixation point contrasts to ensure that the subject only sees the fixation point of the perceived object during the experiment.

### 2.3. Task Definitions

After stimulus calibration, subjects were instructed about the specific requirements of each task that they were going to perform on that day. For each of the four tasks, the session consisted of 16 blocks of 16 trials each. In a single block, there were 8 binocular rivalry (BR) trials, where the subject viewed the overlaid red and cyan objects to experience bistable vision, and 8 “physical” trials where each of the two stimuli was monocularly presented in an alternating fashion (**Fig. 1A**). In tasks that involved shifts in fixation point (FP) locations, new FP locations were defined at the beginning of each trial, out of *four possible fixation point pairs* for the red and cyan FPs. Each nominal trial lasted 5 seconds, but with a random assignment of FP pair locations, we sometimes induced actual trials of 10, 15, or 20 seconds because the new FP locations were the same as in the previous trial. This way, subjects did not habituate to a strict 5-second routine. When we define trials as the number of location shifts of the FPs, subjects underwent on average 91 BR trials (**Supplementary Table 1**).

The different tasks are shown in **Fig. 1B**. In **Task 1** (closest to the original task of Hesse and Tsao (2020)), subjects were instructed to follow the fixation point they are currently perceiving, and press the keyboard button corresponding to the perceived object at the same time. **Task 2** was a continuous binocular rivalry paradigm, where the subjects were instructed to report the perceived object with the corresponding button while maintaining their fixation to the center of the screen. Thus, Task 2 was our negative control task where both tested external factors (shifts in FPs and saccades) were eliminated. Total block length was kept the same as in Task 1 (80 seconds per block, and 16 blocks in total per task). While pre-processing the behavioral data from Task 2, we pseudo-chopped the continuous blocks into trials by defining four possible pseudo-trial states, and the state was chosen randomly at the beginning of each 5 seconds, creating a similar amount of 5, 10, 15, and 20 second trials as in Task 1 for analysis purposes. In **Task 3**, subjects again fixated at the center of the screen with the instruction to ***not*** perform a saccade to the shifting FPs of the objects, but rather to fixate on the central FP while reporting their perception with the keyboard. In **Task 4**, subjects were instructed to perform a saccade from one black FP to another following a brief auditory cue, while reporting their perception with the keyboard. The onsets (times) of these cues were determined by the same rule that determined the onsets of FP shifts in tasks 1 and 3, leading to a similar number of trials. Subjects were told that there may be “transitions” between the two stimuli, where they might see both stimuli at the same time. They were told to release the buttons during transitions or when they perceived parts of both stimuli. In the physical trials of each task, we presented only one of the objects monocularly in each trial, making sure that the subject is engaged in the task by pressing the correct button corresponding to the object presented on the screen.

### 2.4. Eye tracking analysis

Eye tracking data were recorded at 1000 Hz but resampled to 100 Hz for the analysis. The distances between the gaze and the fixation points were computed across time in order to verify that the subjects followed the fixation point of the perceived object, and not the other one.

Although eye tracking results in selected subjects from Task 1 replicated the results from Hesse and Tsao (2020) such as BR perception can be tracked from eye position (**Supplementary Fig. 1**), the close distance between the fixation points (average distance = 3.31°, range of distances = [2.82 3.62]°) was too small for our eye tracking procedures to reliably decode the perceived object from trial-to-trial gaze location data, as performed in Hesse and Tsao (2020). This is likely due to differences in eye tracker systems and calibration procedures, including less rigid head stabilization than in the study of Hesse and Tsao. We do not consider this problematic for our study as our questions rely on task timings and the perceptual reports of the subjects across conditions with and without saccades.

Despite the limitation regarding the continuous gaze tracking for the inference of the perceived object, we could detect saccades performed as part of the task requirements at trial onsets in Task 1 and Task 4. These saccades could be detected reliably in 10 out of 17 subjects due to limited gaze location accuracy. Both the number of subjects that we could use for saccade detection, and the number of saccades that could be detected from some of these subjects were too low to be informative in a group-level analysis where mean or median values from each subject are used (**Table 4**). Therefore, we limit our saccade-locked perceptual switch latency analyses to all saccades pooled across the 10 subjects that we could include (**section 3.4**).

The saccade detection algorithm was adapted from Hesse and Tsao (2020). For an eye movement at time *t* to be detected as saccade in the original algorithm, the distance between the mean gaze location during the last 100 ms and the mean gaze location during the next 100 ms should be larger than 0.5° (change of location criterion), and the gaze location in each time window should stay within 0.5° of the mean gaze location in the respective window (stability criterion). Since the desired saccades for our analyses were the ones at the beginning of the trial, specifically the ones from the old fixation point towards the new fixation point of the perceived object, we used distances to the old and new fixation points instead of x- and y-coordinates of gaze location. Distance criteria were slightly modified such that a gaze shift (a “saccade”) was detected at time *t* if distance to the new fixation point decreased more than 0.5° in the next 100 ms with respect to the previous 100 ms. The stability criterion remained the same. Since the desired saccades are bound to happen soon after the trial onset, we limited the time window of saccade detection to 0-1.5 seconds after trial onsets. Since our saccade detection was aimed at saccades due to fixation point shifts at the trial onset, we restricted the detection to the trials where the perceived object remained the same during the transition from one trial to the next (i.e. excluding the cases in which the subject reported a perceptual switch in the last 100 ms of the previous trial, or the first 100 ms of the next trial).

To compare latencies of saccade-induced perceptual switches to a control task without saccades, we simulated random saccade onsets between 0-1.5 seconds of each binocular rivalry trial in Task 2, by using the *rand()* function in MATLAB. Upper limit of the search window for saccades was chosen as 1.5 seconds because we estimated that it is unlikely for a trial onset-locked saccade would take place after 1.5 seconds from trial onset. For both the real saccades in tasks 1 and 4, and the simulated saccades in Task 2, the first perceptual switch that occurred after the saccade onset in each trial (if any) was used for saccade onset-locked perceptual switch analysis.

### 2.5. Behavioral Instructions and Analysis

Trial onsets were defined as the moment of fixation point (FP) shifts in tasks 1 and 3, and auditory cue onsets in Task 4. Onset of a perceptual switch can be exemplified as follows: Subjects pressed the ALT button on the keyboard when they perceived the red object dominantly. When their perception was a mixture from the two eyes, not enabling them to clearly choose the perceived object, they were told that they do not need to press any button. However, since the mixed percepts sometimes created confusion for the subjects, we saw that there are periods where they pressed both buttons. Following this ‘piecemeal perception’ period, sometimes they went on to press the CTRL button, meaning that they started perceiving the cyan object dominantly. In this case, we categorized such perceptual switch as ‘complete’ and the onset of the switch was the moment when the subject changed the button press from ALT to CTRL+ALT or None (**Fig. 1C**, top row). Sometimes, a period of ‘piecemeal perception’ was followed by a dominant perception of the previous stimulus again (**Fig. 1C**, middle and bottom rows). Such ‘perceptual switches’ were categorized as ‘incomplete’, with the same onset as the ‘complete’ ones. Perceptual Switch Latency (PSL) was defined as the time in between the last trial onset before the perceptual switch, and the onset of the perceptual switch. The number of switches that each subject perceived in each task, and their median latencies are given in **Supplementary Table 1** and **Supplementary Table 2**, respectively. Due to the presence of 10, 15, or 20-second trials, there were rare instances of PSLs longer than 5 seconds. Since they introduce a significant amount of irregularity to the histograms of PSLs and they are not relevant to our main hypotheses, we excluded these rare events from our analyses.

Statistical tests are detailed in the **Results**. However, we followed some general rules, pre- determined before data analysis. We first checked the PSL distributions at the individual and group-level. Since they did not necessarily follow normal distributions (assessed by one-sample Kolmogorov Smirnov test), we chose non-parametric statistical tests for comparisons within and across tasks. All between-task analyses were performed by using a (non-parametric) test for *repeated measures*, since the same set of subjects (n=17) underwent all four tasks. Since the individual latency distributions did not follow normal distributions (one-sample Kolmogorov Smirnov test), we used the median latency values from each individual while performing group-level statistics. In the results, we also report the average (mean) and the standard deviation of these individual median latency values, in the form of “[Mean SD]”. When we applied separate paired-tests to multiple task pairs, which required correction for multiple comparisons, we corrected our default statistical significance threshold (α=0.05) by dividing it by the number of tests in the corresponding analysis (Bonferroni correction). In other cases, where we applied an across-task comparison with one-way Friedman tests, we used the *multcompare()* function in MATLAB for post-hoc comparisons, with Tukey-Cramer correction for multiple comparisons. To compare experimental distributions with theoretically expected uniform distributions, we used *unifrnd()* function in MATLAB to create uniform distributions of the same size as the corresponding experimental distributions.

All scripts regarding the stimulus properties, task implementations, and data analysis can be found in this GitHub repository: https://github.com/dagdpz/BinoRiv_In2PB.

## 3. RESULTS

We hypothesized that two external factors – saccades to new fixation point locations and visual transients due to changing fixation point locations – in the original no-report BR paradigm from Hesse and Tsao (2020), may induce perceptual switches that might not occur in absence of these factors. We thus introduced a two-way factorial experiment to test these hypotheses (**Fig. 1B**). Since our main hypotheses is centered around the question of whether the external factors induce any sort of disruption or change in what the subjects perceive, our statistical analyses focus on the combined (pooling complete and incomplete switches, **Fig. 1C**) perceptual switch latencies (PSLs) with respect to the trial onsets. We provide the number of complete and incomplete perceptual switches of each subject in **Supplementary Table 1**, and the average latencies of these switches in **Supplementary Table 2** and **Supplementary Figures 3** and **4**.

### 3.1. Perceptual Switch Latency (PSL) is influenced by saccade execution

We first compared the median Perceptual Switch Latency (PSL) values between the four tasks, where latency values represent the time between trial onsets (t=0 s; corresponding to external events, namely the fixation point shifts in Task 1 and Task 3, the auditory cue in Task 4, as well as simulated “trial onsets” in the continuous Task 2) and the report of perceptual switch by the subject (**Fig. 2B**). As shown in **Fig. 2B**, the PSL of Task 1, where saccade and FP jump co-occur together and the PSL of task 4 that included saccades only, look shorter than the other tasks. Accordingly, we found that task identity had a main effect on PSLs (n=17, one-way Friedman’s test: χ^2^=20.15; p=1.58x10^-4^). Post-hoc comparison of PSLs between task pairs revealed that PSLs between task pairs 1-2 and 2-4 differed significantly, with Task 1 and Task 4 having significantly lower median PSLs than Task 2 (**Fig. 2B** and **Table 1**). We also grouped the tasks according to the presence of saccades (tasks 1&4 vs. tasks 2&3) and shifting fixation points (tasks 1&3 vs. tasks 2&4), and compared the median PSLs between these groups. Two-way repeated measures Friedman’s test (task identity is also taken into account as a grouping factor) revealed that *saccade requirement* had a main effect on PSLs (χ^2^ = 12.99, p=3.12x10^-4^, **Fig. 2C**), whereas *shifts in fixation point locations* did not (χ^2^ = 0.21, p=0.64, **Fig. 2D**). These results show that presence of external events can influence the time course of perceptual switches in this BR paradigm, and some external factors (saccade requirement) may have a stronger influence than other factors (binocular visual transients introduced by shifting fixation points).

**Figure 2:**
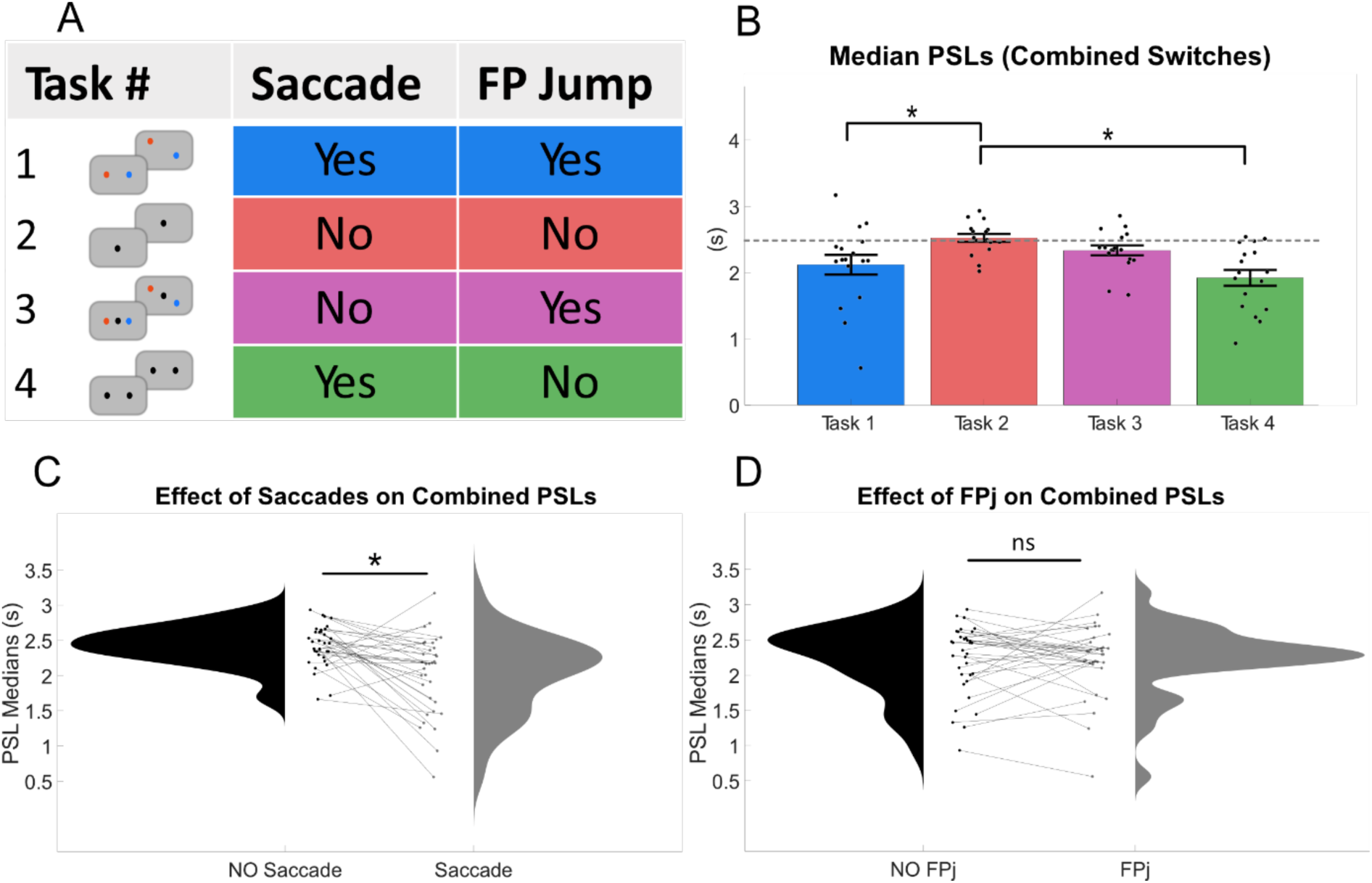
Median Perceptual Switch Latency in each task. **(A)** Categorization of tasks based on the presence of saccade requirements and alternating fixation points in each trial. **(B)** Bar graphs show the group-level average perceptual switch latencies (PSLs), pooling the complete and incomplete perceptual switches. Each data point represents the median PSL of one subject. Type of task had a significant main effect on PSLs, and post-hoc tests revealed significant differences between PSLs in tasks 1 vs. 2, and tasks 2 vs. 4 (shown with *). **(C)** and **(D)** PSLs after grouping the tasks according to the presence of saccade and the presence of fixation point jump (FPj). Connecting lines represent single subjects. Friedman’s test, adopted as the non-parametric version of two-way repeated measures ANOVA for this particular analysis, revealed a main effect of saccade but no main effect of jumping fixation points.

**Table 1:** Summary of statistical analyses regarding Perceptual Switch Latencies (PSLs) across tasks (corresponding to Fig. 2B). Left 3 columns: group-level mean of the median PSLs. Right 5 columns: post-hoc pairwise comparison of PSLs. P-values correspond to corrected values for multiple comparisons.

| COMBINED PSLs - All Switches |  |  |  |  |  |  |  |
| --- | --- | --- | --- | --- | --- | --- | --- |
| Single Task Average Values |  |  | Post-hoc pairwise comparison of PSLs |  |  |  |  |
| Task | Mean | SD | Task A | Task B | Diff | p | Confidence Interval (95%) |
| 1 | 2.12 | 0.61 | 1 | 2 | -1.41 | **0.01** | [-2.55 -0.27] |
| 2 | 2.52 | 0.24 | 1 | 3 | -0.59 | 0.54 | [-1.73 0.55] |
| 3 | 2.34 | 0.31 | 1 | 4 | 0.47 | 0.71 | [-0.67 1.61] |
| 4 | 1.92 | 0.49 | 2 | 3 | 0.82 | 0.25 | [-0.31 1.96] |
|  |  |  | 2 | 4 | 1.88 | **1.25E-04** | [0.74 3.02] |
|  |  |  | 3 | 4 | 1.06 | 0.08 | [-0.08 2.20] |

In the absence of an external event that induces a time-locking effect, the median point of all switch events should converge to 2.5 seconds, the middle of a 5-second trial (see **Supplementary Fig. 2** for simulations, which revealed a median simulated PSL value of 2.50 seconds in the absence of any external events occurring at trial onsets). We tested the median PSL values from each task against this theoretically expected value (**Fig. 3**, left). We found that median PSLs in Task 1 ([Mean SD] = [2.12 0.61]) and Task 4 ([Mean SD] = [1.92 0.49]) were significantly closer to the trial onsets than 2.5 seconds (one sample Wilcoxon signed-rank test against 2.5 seconds; p=6.49x10^-3^ and 4.55x10^-4^, respectively). In contrast, latencies in Task 2 ([Mean SD] = [2.52 0.24]) and Task 3 ([Mean SD] = [2.34 0.31]) were not (p = 0.59 and 0.03, respectively). The p-value corresponding to Task 3 shows a trend towards a time-locking effect, but does not survive our correction for multiple comparisons (α = 0.05/4 = 0.0125). We then compared PSLs pooled across all subjects with a uniform distribution of the same size in each task (generated by unifrnd() function in MATLAB), since theoretically a perceptual switch can happen anywhere within a trial in the absence of any external factors, resulting in a uniform distribution of PSLs (**Fig. 3**, right; see also **Supplementary Fig. 2C**). We found that PSL distributions in Task 1 and Task 4 differ significantly from a uniform distribution (two-sample Kolmogorov-Smirnov tests; p = 7.80x10^-8^ and 1.03x10^-11^, respectively) and it is visible on the histograms that perceptual switches occur more frequently at times closer to trial onsets. Conversely, PSL distributions in Task 2 and Task 3 were not significantly different from the corresponding uniform distribution (p = 0.98 and 0.14, respectively).

**Figure 3:**
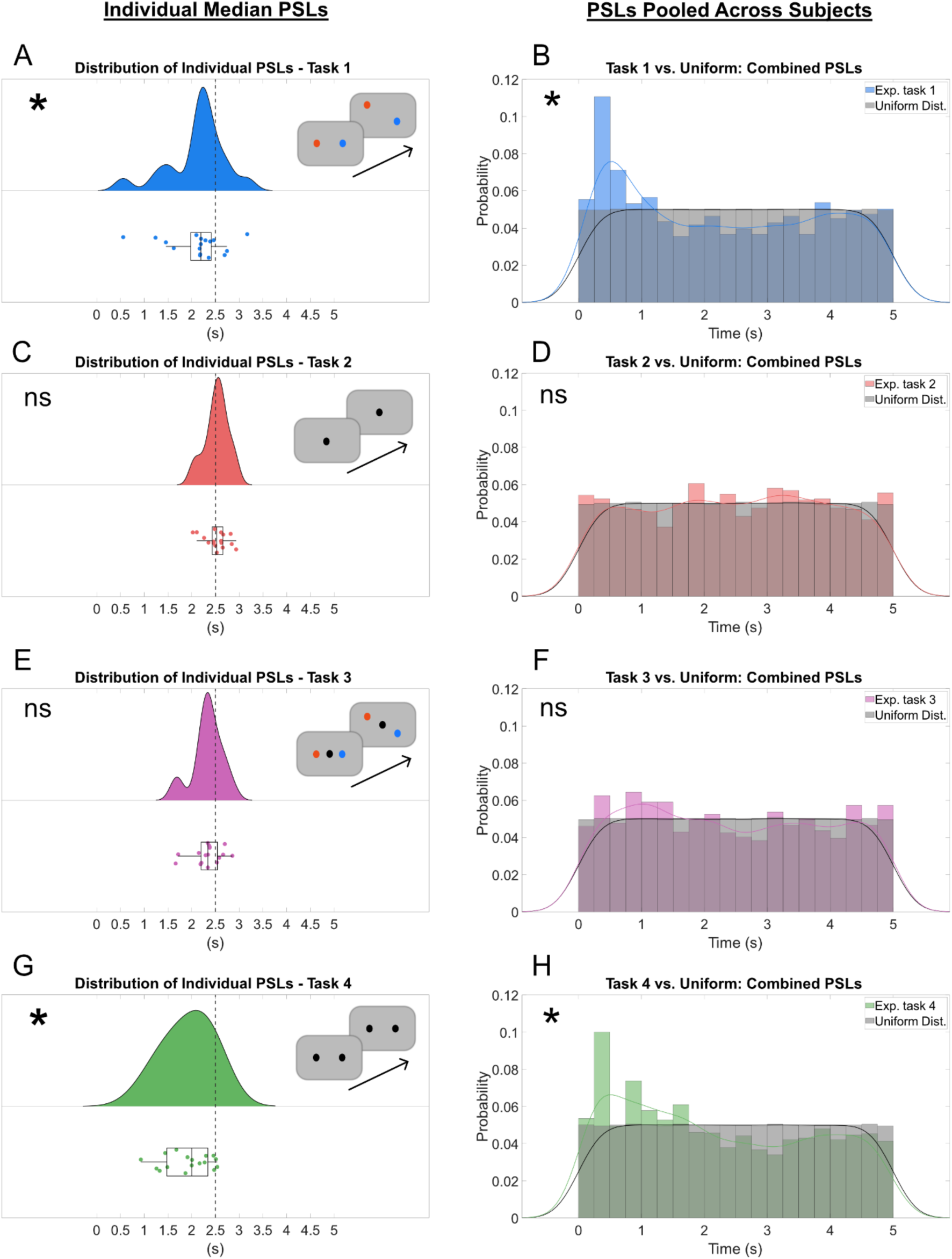
Median and pooled PSLs as a function of task. Median PSLs from each individual constituted a group-level distribution with a significantly smaller (denoted by *) median value than 2.5 in tasks 1 and 4 **(A, G)**, but the medians of the corresponding distributions were not significantly smaller than 2.5 in tasks 2 and 3 **(C, E)**. **(B, D, F, H)** We also compared the perceptual switch probability distributions from each task with a uniform distribution of the same sample size as the experimental distributions. As expected, switch latency distributions from Task 2 did not differ from the uniform distribution. We also did not see a significant difference in Task 3, despite a trend towards a higher likelihood of perceptual switches closer to trial onsets. In contrast, we found significant time-locking effects in Task 1 and 4. Solid lines are fitted curves for visualization purposes. Distributions were compared by Kolmogorov-Smirnov tests (*: p < 0.05 after correction for multiple (four) comparisons).

So far, we considered all switches in a trial, regardless whether they were complete or incomplete. We thus performed an additional analysis where we consider only complete switches (**Supplementary Fig. 3**). Distribution of median PSLs from each subject did not show a significant difference from the theoretically expected average latency value (2.5 seconds), but the experimental average latencies still showed a trend towards shorter latencies than 2.5 seconds (i.e., a trend towards time-locking to the trial onset). The general result that the switch latencies are time-locked to trial onsets in the tasks with saccade condition remained, based on the distributions of PSLs pooled across subjects (**Supplementary Fig. 2,** section SR. 1).

Similar analysis by using only incomplete perceptual switches shows that they are more likely to occur shortly after trial onsets in the tasks with saccade requirements (**Supplementary Fig. 4**), while also revealing that subjects experience significantly more incomplete perceptual switches in the original paradigm from Hesse and Tsao (Task 1), with respect to our experimental negative control (Task 2) (**Supplementary Results,** section SR.1).

Since Task 2 was specifically designed as the experimental negative control with no saccade requirement and no changes in fixation point locations, we compared median PSLs in each task to the median PSLs in Task 2 (**Fig. 4**, left column). Statistical results are similar to what is shown in **Fig. 2B** (Wilcoxon signed-rank test (n=17); Task 1 vs. Task 2: p=0.01, Task 3 vs. Task 2: p=0.16, Task 4 vs. Task 2: p=8.46x10^-4^). In addition, we compared the distribution of PSLs pooled across all subjects and compared each distribution to Task 2. Results were similar such that PSL distributions in tasks 1 and 4 significantly differed from that of Task 2 (p=6.05x10^-9^ and p=1.36x10^-11^, respectively, **Fig. 4**, right column). In addition, PSL distributions between Task 3 and 2 also differed significantly (p=0.01, **Fig. 4D**).

**Figure 4:**
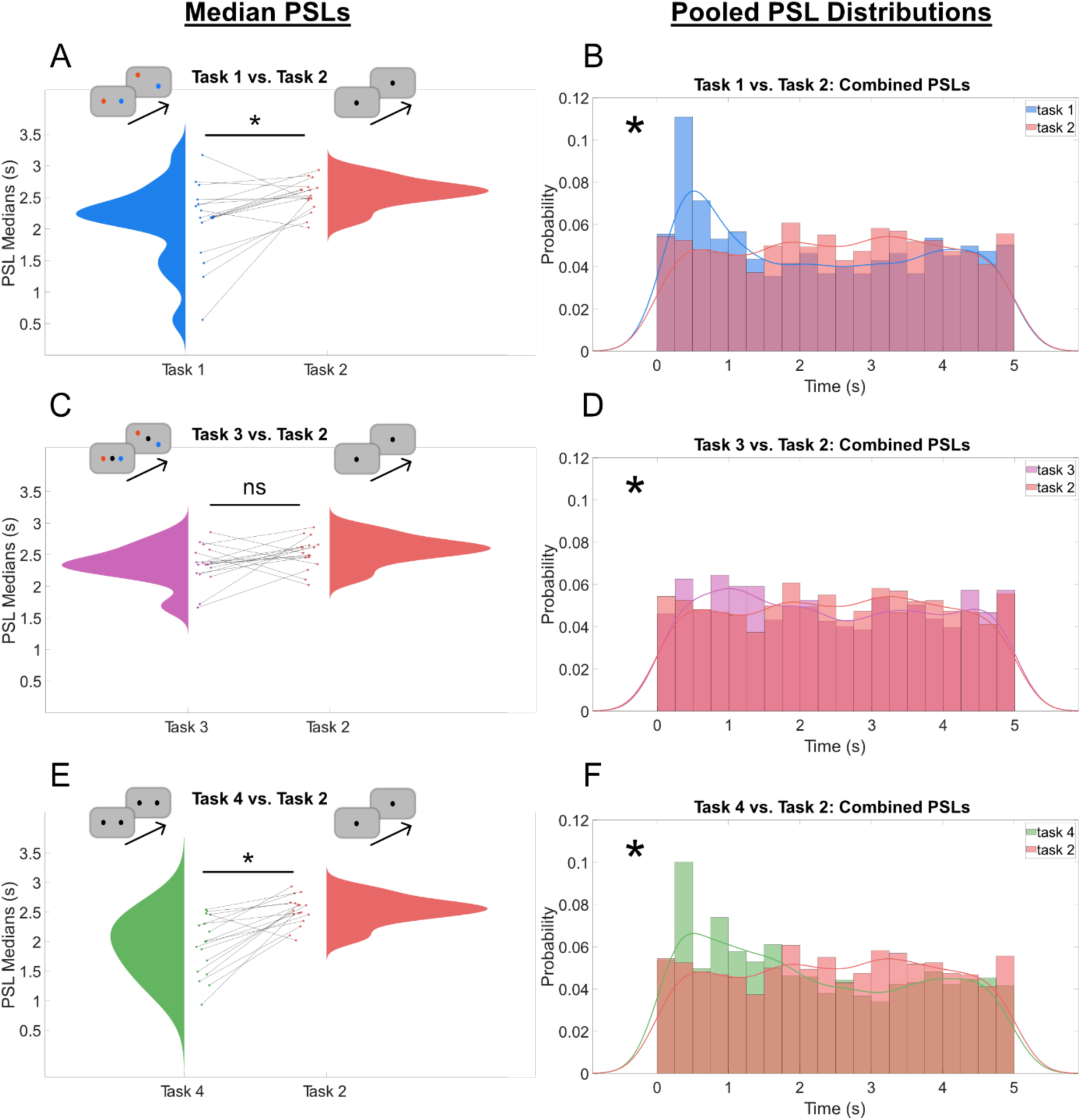
**Task-pair comparisons of median and pooled PSLs at the group-level**. Each task is compared against Task 2 since this task has been designed as our experimental negative control. PSLs in tasks 1 **(A-B)**, and 4 **(E-F)** are found to be significantly more time-locked to the trial onset in comparison to Task 2 (*: p<0.05 after correction for multiple comparisons, dividing the raw p-value by the number of comparisons – three – in this case). Medians of combined perceptual switch timings from each individual are shown as scattered data in **A**, **C**, and **E**, connecting lines represent single subjects. Corresponding group-level comparisons are performed with non-parametric paired Wilcoxon signed-rank tests. Histograms **(B, D,** and **F)** are obtained after pooling perceptual switch latencies across individuals. Comparisons of distributions are performed with two-sample Kolmogorov-Smirnov tests (*: p<0.05 after correction). Subject 3 was excluded from comparative analysis with Task 2 because this subject underwent much fewer trials in Task 2 than any other subject (**Supplementary Table 1**).

### 3.2. PSL Analysis for the *First* Perceptual Switches

So far, we have considered all perceptual switch events, irrespective of their timings within the trial. We showed that perceptual switches are more likely to occur at the beginning of trials due to fixation point shifts and subsequent saccades (**Figs. 2-4**, see **Supplementary Fig. 3** for completed switches only). However, the distributions of perceptual dominance durations show that the time between most of the successive perceptual switches is less than 3-4 seconds (**Supplementary Fig. 5A-F**). As a result, some of the perceptual switches that occur right after the trial onset (i.e., within the first second of a trial), are followed by another perceptual switch towards the end of the 5-second period (**Supplementary Fig. 5G-H**). In order to further isolate the potential time-locking of perceptual switches to external factors, we selected the switch events that are the *first switches within their respective trials* and excluded further (secondary) perceptual switches. The choice of this subset expectedly skews the median PSLs closer to trial onsets even in Task 2, because excluding secondary switches means that we bias switch events towards the beginning of the trial. Comparing the timings of first switches across tasks (**Fig. 5B**), we found a main effect of task type on first PSLs within trials (one-way Friedman’s test; χ^2^ = 19.38, p=2.29x10^-4^). Post-hoc tests with correction for multiple comparisons revealed that the medians of first PSLs differ significantly between the task pairs 1-2, 2-4, and 3-4 (**Fig. 5B** and **Table 2**). Possibly the comparison between tasks 1 and 3 did not reach significance (p=0.06) as the visual onsets of the new fixation spots also had a small effect on switch probabilities. Two-way repeated measures Friedman’s test revealed that saccades have a main effect on the latencies of first perceptual switches (χ^2^=21.64, p=3.29x10^-6^, **Fig. 5C**), whereas shifts in fixation point locations does not (χ^2^=0.43, p=0.51, **Fig. 5D**). This result is consistent with the analysis where we considered all perceptual switches.

**Figure 5:**
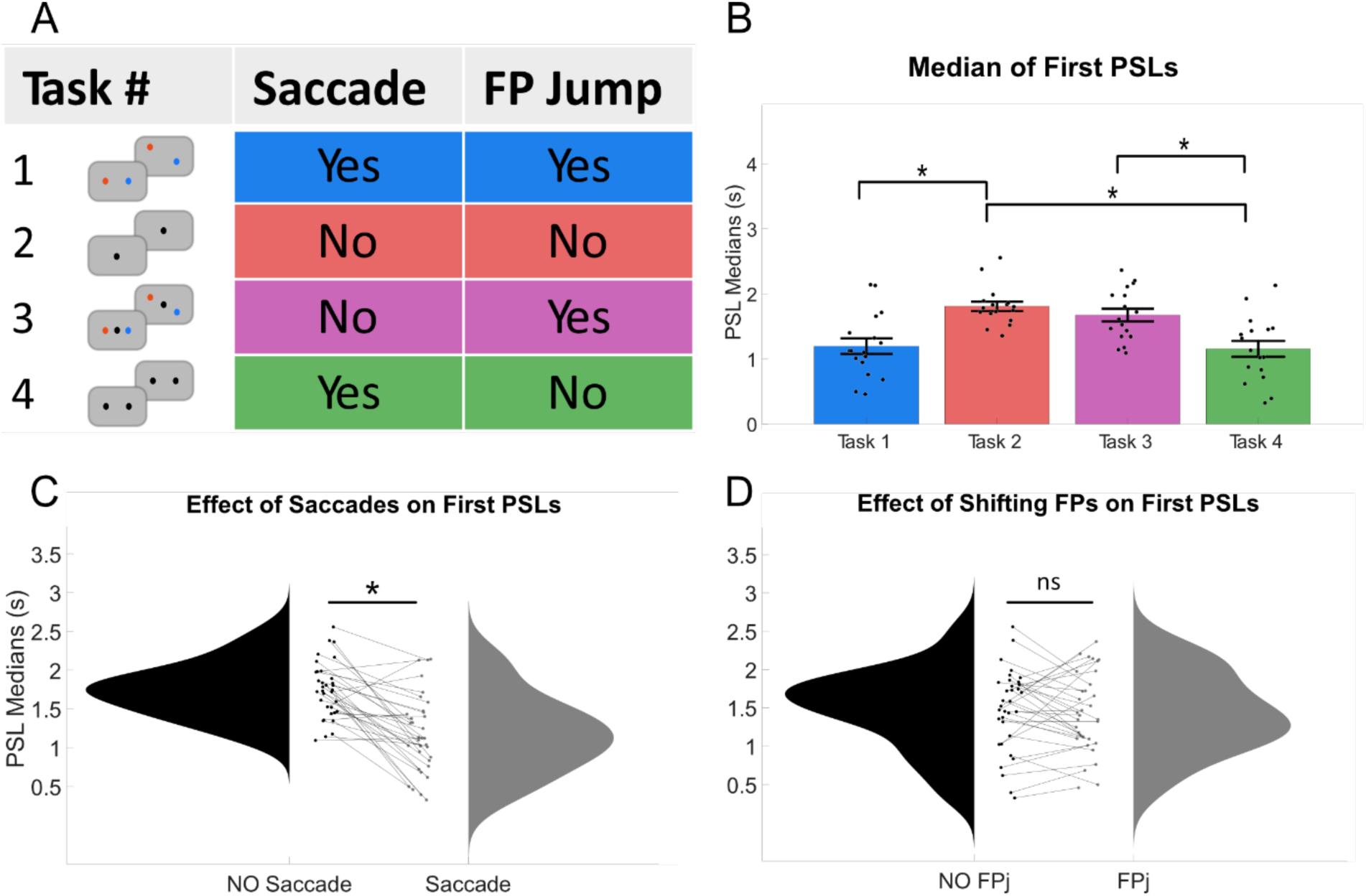
Median PSL comparisons across tasks 1-4, while using only the *first* perceptual switch within trials, excluding secondary events. Here, we have no theoretical expectations as to where the median PSLs should be in the absence of external factors, since we only select a subset of all switches. **(A)** Categorization of tasks based on the presence of saccade requirements and alternating fixation points in each trial. Across tasks, first PSL comparisons reveal that task identity has a significant effect on the latencies of first perceptual switches. **(B)** Task pairs that significantly differ from each other after correction for multiple comparisons are shown with asterisks. When grouped according to the presence of saccades **(C)** and jumping fixation points **(D)**, we found a main effect of saccades on the first PSLs, and no effect of shifts in fixation point locations.

**Table 2:**
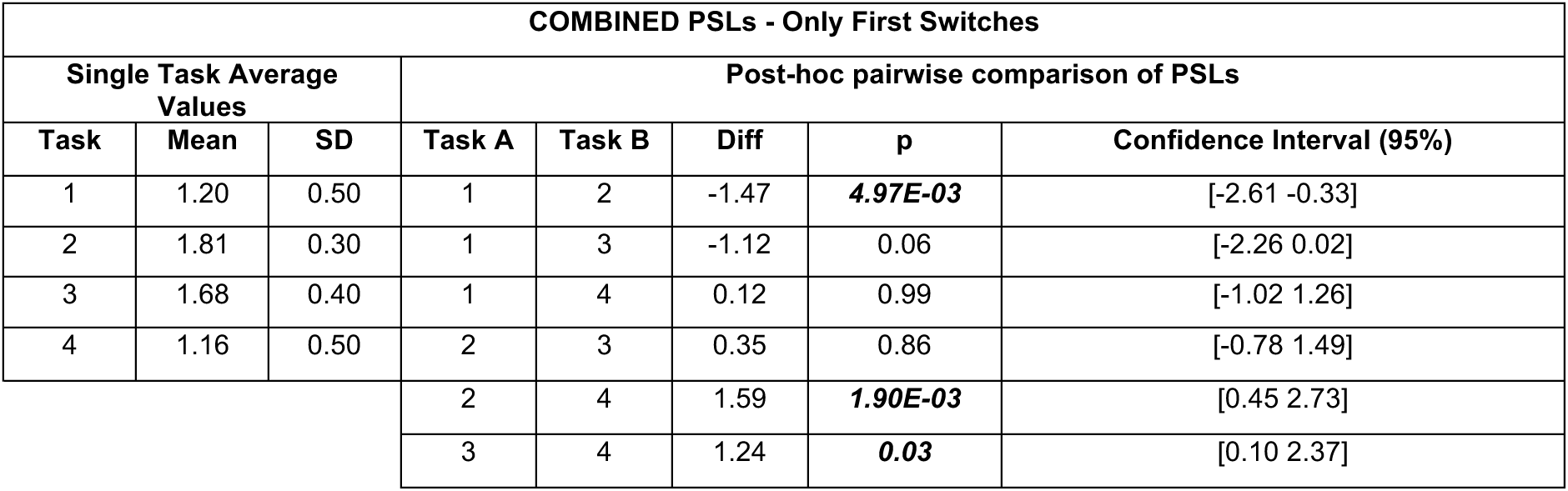
Summary of statistical analyses across tasks regarding first perceptual switch latencies. Left 3 columns: group-level mean of the median PSLs when only the first perceptual switches within trials are used. Right 5 columns: post-hoc pairwise comparison of PSLs. P-values correspond to corrected values for multiple comparisons.

| COMBINED PSLs - Only First Switches |  |  |  |  |  |  |  |
| --- | --- | --- | --- | --- | --- | --- | --- |
| Single Task Average Values |  |  | Post-hoc pairwise comparison of PSLs |  |  |  |  |
| Task | Mean | SD | Task A | Task B | Diff | p | Confidence Interval (95%) |
| 1 | 1.20 | 0.50 | 1 | 2 | -1.47 | **4.97E-03** | [-2.61 -0.33] |
| 2 | 1.81 | 0.30 | 1 | 3 | -1.12 | 0.06 | [-2.26 0.02] |
| 3 | 1.68 | 0.40 | 1 | 4 | 0.12 | 0.99 | [-1.02 1.26] |
| 4 | 1.16 | 0.50 | 2 | 3 | 0.35 | 0.86 | [-0.78 1.49] |
|  |  |  | 2 | 4 | 1.59 | **1.90E-03** | [0.45 2.73] |
|  |  |  | 3 | 4 | 1.24 | **0.03** | [0.10 2.37] |

We also compared the group-level median PSL distributions in each task with that of Task 2, using only the first switch events within trials, excluding the secondary events (**Fig. 6**). Results are similar to the respective analysis we performed when we included the secondary perceptual switch events (see **Fig. 4**), but the differences between the median PSL distributions of *only the first perceptual switches within trials* are more pronounced. Again, we found that Task 1 and Task 4 have significantly smaller median PSLs at the group level (Wilcoxon signed-rank test; p=8.46x10^-4^, and p=3.52x10^-4^, respectively). Median PSLs of the first switches did not significantly differ between Task 2 and Task 3 (p=0.04, trend fails to reach statistical significance after correction for multiple comparisons) (**Fig. 6**, left column). On the other hand, pooled distribution of first PSLs in each of tasks 1, 3, and 4 differed significantly from that of Task 2 (p=2.63x10^-15^, p=6.97x10^-4^, p=3.84x10^-25^, respectively; **Fig. 6**, right column). Assessment of the p-values again reveals that the effects of external factors in Task 1 and Task 4 (saccades) are larger when compared to that of Task 3 (visual transients due to fixation point shifts), which is also confirmed via the post-hoc tests shown in **Fig. 5B** and **Table 2**.

**Figure 6:**
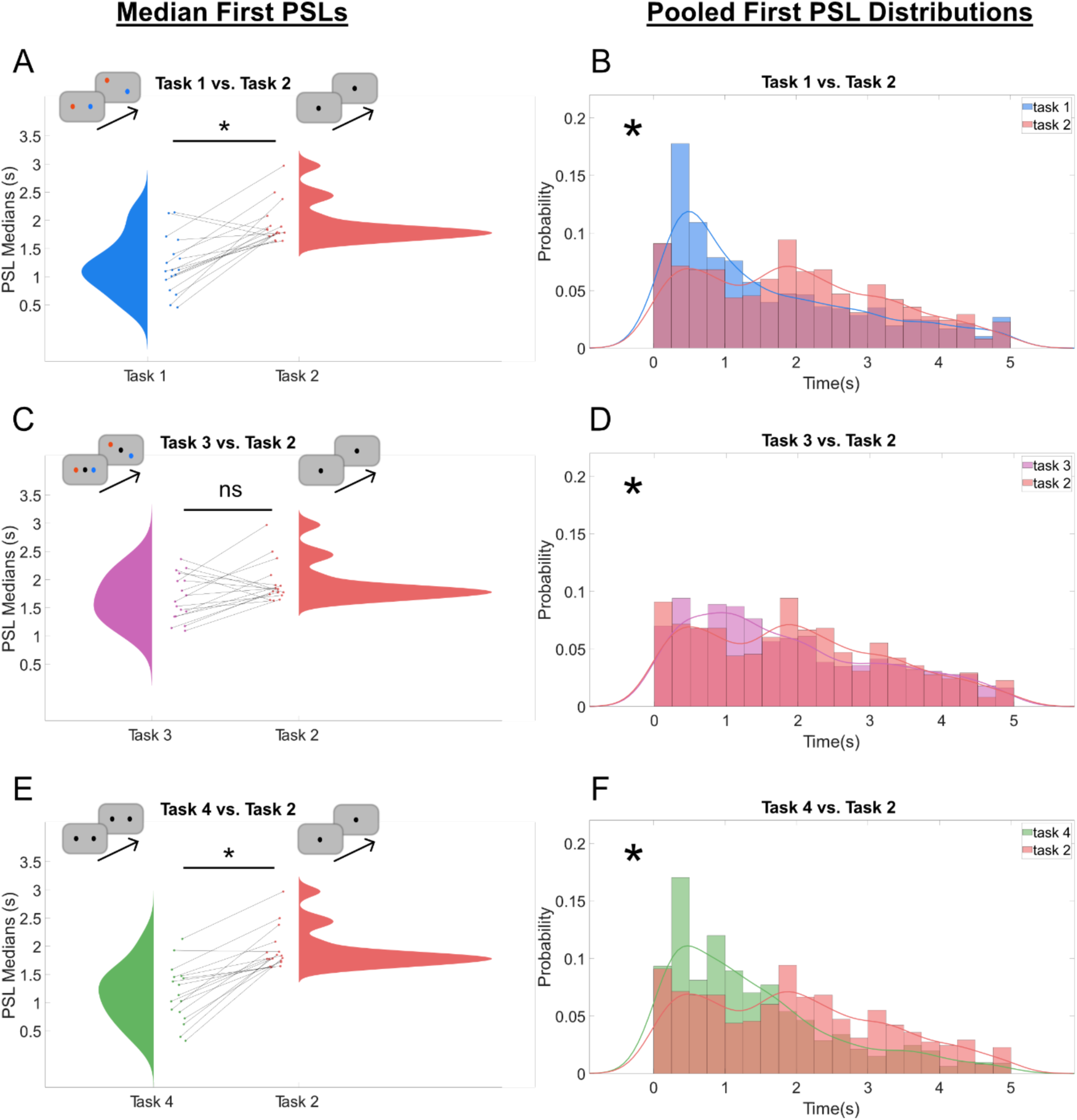
**Task-pair comparisons of PSLs *of only the first switch events*** (differing from the analysis in Fig. 4 in this regard) within trials at the group-level. Each task is compared against Task 2. Connecting lines in **A, B** and **E** represent single subjects. PSLs of *first* switches in tasks 1 **(A-B)**, and 4 **(E-F)** are found to be significantly more time-locked to the trial onset in comparison to Task 2 (*: p<0.05 after Bonferroni correction for three comparisons). Medians of the first PSLs in Task 3 are not significantly different from Task 2 **(C)**, but the PSL distributions of tasks 2 and 3 pooled across subjects are found to differ significantly **(D)**.

*Median* point of a distribution may not always accurately represent the data points with the maximum likelihood (e.g., a symmetrical U-shaped distribution would have the middle data point as the median, whereas the data points with the maximum likelihood would be the left- and right-ends of the distribution). Thus, we performed an additional analysis on the *maximum latency probabilities* in each task, instead of using the median values (**Supplementary Results**, section SR.3). Similar to the previous analyses, grouping the tasks according to the presence of saccade requirements and shifting fixation points showed that the saccade requirement has a main effect on PSL (**Supplementary Fig. 8**). In addition, the latency bin with the maximum probability is significantly closer to trial onsets than the theoretically expected value (2.5 s) in the original paradigm (Task 1) but not the other tasks (**Supplementary Fig. 9A-D**). When compared to the experimental control (Task 2), PSLs with the maximum probability in tasks 1 and 4 are significantly closer to the trial onsets (**Supplementary Fig. 9E-G**), supporting the results obtained from the median PSLs (**Fig. 4**).

### 3.3. Can the spontaneous and induced perceptual switch events be separated in this no-report binocular rivalry paradigm?

In order to investigate the neural mechanisms behind **spontaneous** changes between perceptual states, future studies may find it useful to select a subset of perceptual switch events that are not time-locked to an external event. We thus calculated the number of *complete* perceptual switch events that occurred after the *first* perceptual switch and before another trial onset (*secondary* complete perceptual switches). In Task 1, 35% of all *complete* perceptual switches pooled across subjects are *secondary* perceptual switches (a total of 464 events out of 1327 *complete* perceptual switches). These events are probably independent from FP shifts or saccade planning/execution at the trial onset since another perceptual switch takes place before the *secondary* event. When we look at the secondary switch ratios in each subject, we see that 32.3% of all complete perceptual switches are *secondary* switches (median across subjects), and thus usable in the analyses of spontaneous changes between perceptual states (**Table 3**).

**Table 3:**
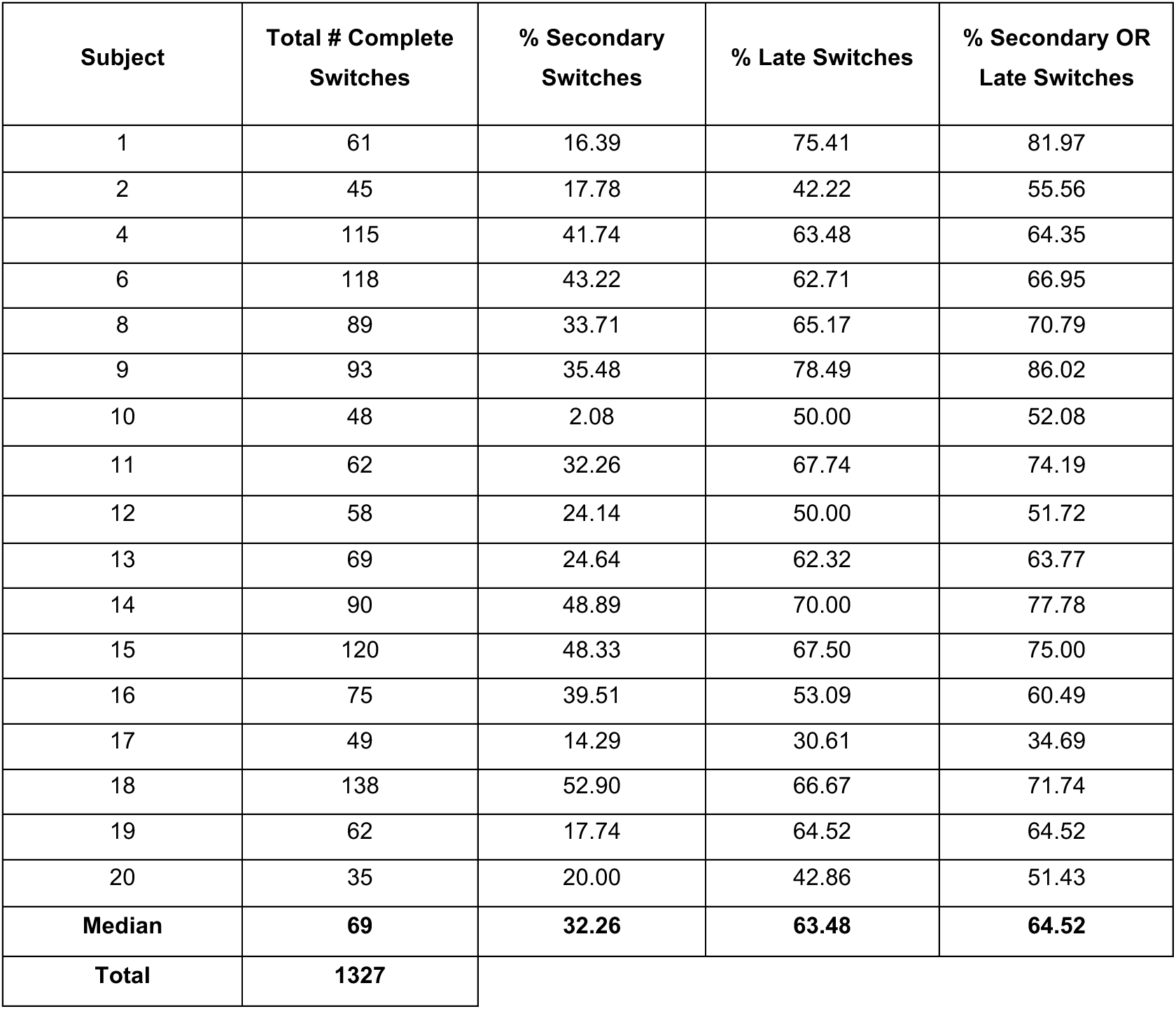
The number of complete switches, and the percentage of complete switches that are secondary or “late”.

A less conservative approach to extract spontaneous perceptual switches would be to estimate the time window in which external triggers (i.e. trial onsets in Task 1) induce a perceptual switch with a higher probability than the pseudo-trial onsets in Task 2. As seen in **Fig. 6B**, such a difference is present in the first 1.5 seconds following the trial onsets, with perceptual switch probability visibly higher in Task 1 than Task 2. We calculated the ratio of complete perceptual switches with a latency larger than 1.5 seconds for each subject (**Table 3**). In Task 1, 62.6% of all *complete* perceptual switches pooled across subjects are “*late”* perceptual switches (a total of 830 events out of 1327 *complete* perceptual switches). The most flexible approach would be to combine the first two approaches. One could say that a perceptual switch is independent of the external factors if it is at least 1.5 seconds later than the trial onset, *or* it is a secondary perceptual switch. In our data, 67.8% of all perceptual switches satisfied these criteria (899 events out of 1327). When we look at the ratio of perceptual switches that are “late” or secondary in each subject, we see that a median of 64.5% of perceptual switches could be categorized as spontaneous switches.

### 3.4 Saccade-Locked Perceptual Switch Analysis

So far, we established that the presence of an eye position change requirement at trial onsets (fixate new FP position, which is typically implemented via saccade) induced significantly more perceptual switches early in the trials, strongly suggesting that a considerable fraction of perceptual switches in these tasks is induced by these external factors. After finding that saccades had a larger influence than the abrupt fixation point shifts on time-locked perceptual switches (see **Fig. 2C-D** and **Fig. 5C-D**), we decided to analyze perceptual switch latencies time-locked to the saccade onsets (as detected from eye-tracking data) instead of the trial onsets. The reasons for conducting this additional analysis are two-fold. Firstly, knowing the saccade onset time could lead to more accurate estimation of timing of externally-triggered perceptual switches. Secondly, subjects might not have fully complied with the task instructions and did not always fixate the new FP position, which could lead to underestimation of externally-triggered effects.

Due to the limited eye-tracking accuracy (see **Supplementary Fig. 1**), only 10 out of 17 subjects could be used for these saccade onset-locked analyses, and only the saccades at trial onsets could be reliably detected (see **Methods**). Numbers of detected saccades (Task 1 and Task 4) or simulated saccades (Task 2), and numbers of trials with a perceptual switch that follows the detected saccade are shown in **Supplementary Table 4**. Since the number of such perceptual switches per subject that we could analyze from Task 1 and Task 4 are relatively low, we mostly focus on pooled perceptual switches across subjects below. For this analysis, we specifically targeted the saccades that occur at the beginning of trials towards the new location of the fixation point of the perceived object (**Fig. 7A**), because we wanted to quantify the effect of task-related saccade onsets on the perceptual switches, instead of simply time-locking our PSL analysis to the onsets of the trials (and assuming that each fixation point shift is followed by a saccade).

**Figure 7:**
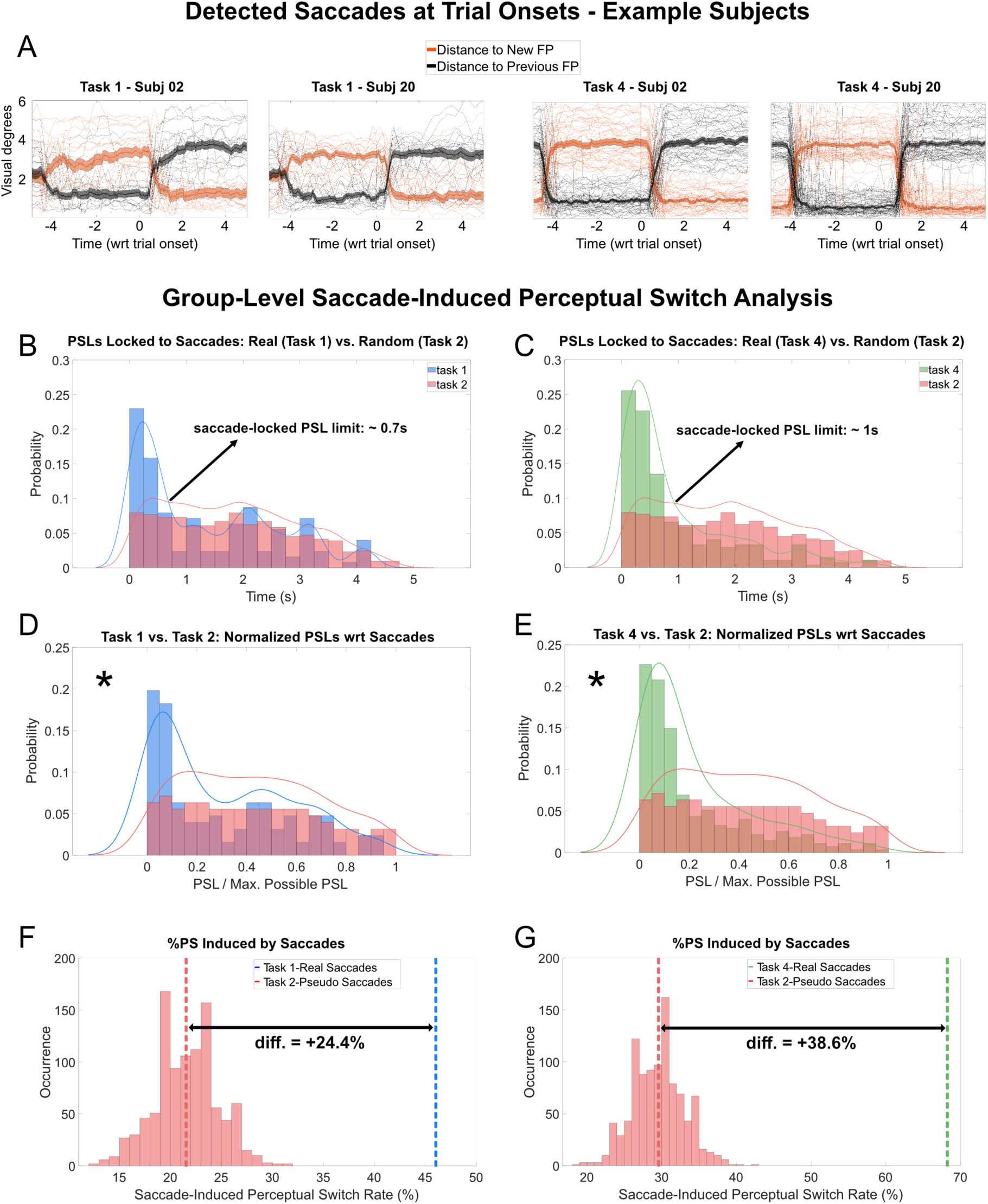
Saccade-locked perceptual switch analysis. **(A)** Single-subject examples (subject #02 and #20) of traces that are detected as a saccade: first two columns are from Task 1 and the second two columns are from Task 4. **(B-C)** Probability distributions of PSLs that follow a detected or a pseudo-saccade. Maximum latency to expect saccade-locked perceptual switches are estimated as 0.7 seconds and 1 second for Task 1 and Task 4, respectively. **(D-E)** Probability distributions of saccade-locked and normalized PSLs show that both Task 1 and Task 4 have a significantly different distribution than Task 2, caused by the increased probability of early perceptual switches in both tasks. Asterisks denote statistically significant difference between the histograms. **(F-G)** Comparing the real (tasks 1 and 4) vs simulated (Task 2) saccade-related switch rates, 24.4% and 38.6% of perceptual switches are estimated to be caused by a preceding saccade in Task 1 and Task 4, respectively.

We first obtained histograms of the latencies corresponding to the first perceptual switches that occur after a detected saccade (or simulated saccades in Task 2). As expected, the probability of early perceptual switches following the real saccades in tasks 1 and 4 is higher than in Task 2, confirming that saccades induce perceptual switches in these tasks (**Fig. 7B-C**). In order to estimate the time window in which a perceptual switch is likely to be induced in each of these tasks, we computed estimated kernel densities of the probability distributions and obtained the time points after which the probability of a perceptual switch in Task 1 or Task 4 is not higher than in the negative control Task 2 (**Fig. 7B-C**: continuous curves). This latency was estimated as 0.70 seconds for Task 1, and 0.97 seconds for Task 4. Since these are qualitative estimates and their exact values depend on the bandwidth of the density estimates, we rounded the latency border of Task 4 to 1.0 second for simplicity. We found that 58 out of 227 detected saccades are followed by a perceptual switch within 0.7 seconds in Task 1, leading to a ratio of 25.5%. The same ratio is only 11.2% for Task 2, meaning that 14.3% of saccades are estimated to “cause” a perceptual switch in Task 1. Similarly, 39.5% of detected saccades are followed by a perceptual switch within 1 second in Task 4, while the same fraction is 15.4% in Task 2, meaning that 24.1% of saccades are estimated to “cause” perceptual switches in Task 4 (**Supplementary Table 3**). We also compared the fraction of saccades that “cause” a perceptual switch by paired t-tests at the group-level (using each subject’s ratio of induced saccades in each task). Even though the numbers of saccades (therefore also the saccades that induce a perceptual switch) that we could detect *per subject* were quite low, we still found that Task 1 and Task 4 have significantly higher ratios of saccades that induce a perceptual switch than Task 2 (t(9): 2.89, p=0.02; and t(9): 4.70, p=1.12x10^-3^, respectively).

Latency of perceptual switches with respect to a preceding saccade can take different possible maximum values in each trial, depending on the saccadic reaction times relative to the trial onset: Task 1 [0.56 0.27] seconds, Task 4 [0.55 0.28] seconds ([Mean SD]). For instance, if a saccade occurs 0.5 seconds after the trial onset, the following perceptual switch can be at most 4.5 seconds later in a 5-second trial; whereas if the saccade has 1 second latency, the following perceptual switch can occur at most 4 seconds later. Therefore, we normalized the saccade-locked PSLs with respect to the maximum value that each latency can get, and compared probability densities between Task 1 and Task 2 (**Fig. 7D**), and between Task 4 and Task 2 (**Fig. 7E**). Both Task 1 and Task 4 had significantly different PSL distributions compared to Task 2, as tested by two-sample Kolmogorov-Smirnov tests, p=1.52x10^-4^ and p=1.32x10^-13^, respectively).

Lastly, we estimated the fraction of perceptual switches that are putatively "caused” by a saccade (ratio of switches that occur within 0.7 seconds for Task 1, and within 1 second for Task 4). Since we simulated a pseudo-saccade onset within initial part of each trial in Task 2 (0-1.5 seconds as a permissive estimate of a trial onset-locked timing), the number of pseudo-saccades is much higher in this task than the detected (real) saccades in tasks 1 and 4 (**Table 4**). As a result, the sample of saccade-locked perceptual switch latencies are also much larger. Thus, we implemented a non-parametric permutation test where we choose random subsets of pseudo-saccade-locked PSLs of the same size as Task 1 or Task 4 in each iteration (n=1000), and calculated the fraction of perceptual switches that occur in the time windows identified above (*cf.* **Fig. 7B,C)**. We found that all pseudo-saccade-locked perceptual switch rates are lower than the real-saccade-locked perceptual switch rates in tasks 1 and 4 (**Fig. 7F-G**). Comparing the putatively saccade-induced perceptual switch rate in Task 1 and the average rate of pseudo-saccade-induced perceptual switches in Task 2, we found that real saccades in Task 1 increased the percentage of saccade-locked perceptual switches by 24.4% (**Fig. 7F**). A similar comparison between Task 4 and Task 2 yielded a 38.6% increase in the percentage of saccade-locked perceptual switches in Task 4. These two values (24.4% and 38.6%) can be summarized as the ratio of post-saccadic perceptual switches “caused” by the real saccades in the corresponding task.

## 4. DISCUSSION

Here, we investigated the perceptual switch latency (PSL) distributions in four variations of a novel binocular rivalry paradigm from Hesse and Tsao (2020), where each rivaling stimulus has a fixation point that changes location in every trial, and subjects are asked to follow the fixation point they see. We found that when the subjects are required to perform a saccade at the onset of each trial (either due to shifts in FP locations in Task 1, or due to the auditory cue in Task 4), the perceptual switch likelihood is significantly increased after these events. The shift in the FP locations without associated saccade has a milder effect on inducing perceptual switches (as tested with Task 3), reaching statistical significance in some of the analyses while staying at the level of a non-significant trend in others.

### Contributions of saccades to perceptual switches during binocular rivalry

There has been a long debate whether eye movements play an important role in perceptual switches. Indeed, it has become clear that when binocular rivalry is stabilized on the retina, e.g. by using afterimages, perceptual reversals still occur (Blake et al., 1971), but the typical dominance duration is much longer compared to normal rivalry conditions (Wade, 1974). Thus, while saccades are not necessary for inducing perceptual switches, they seem to be facilitators. In one study that directly addressed the interaction between perceptual reversals and saccadic eye movements under natural viewing and voluntary percept holding conditions, the authors found a higher saccade probability before the switch in both conditions (Van Dam and Van Ee, 2006). The same study also found that when subjects are asked to try holding one stimulus (e.g., house) as their dominant percept versus holding the other stimulus (e.g., face), they fixate on different locations on the screen, suggesting that the location of fixation might play a role in determining the dominant percept in these paradigms. In a further study that directly addressed the influence of saccade-caused retina image shifts and binocular rivalry reversals, the authors found that the saccade should be accompanied by a shift in the retinal image to induce a perceptual switch (Van Dam and Van Ee, 2006). Similarly, Ross & Ma-Wyatt found that when binocular rivalry stimuli were shown intermittently, saccade in the blank intervals shortened the perceptual durations (Ross & Ma-Wyatt, 2004). Thus, the time-locked perceptual switch effect to the instructed saccade, and the significant increase in perceptual switch probability right after the saccade onsets that we observed in our study, are in line with this previous literature.

### Contributions of shifts in fixation point locations to perceptual switches

One aspect of the Hesse and Tsao paradigm is that the fixation points change their position (i.e., causing visual input to both eyes to shift). Numerous studies probed the impact of short stimulus flashes in one eye on perceptual thresholds and perceptual reversals (Blake et al., 1990; Fox & Check, 1968; Metzger & Beck, 2020). Intuitively, detection thresholds for flashes presented to the dominant eye are much lower compared to the suppressed eye. Importantly, a flash presented to the suppressed eye can force it to become dominant, albeit the resulting individual dominance time is typically shorter (Blake et al., 1990). Importantly, even small stimuli presented to the suppressed eye tend to induce a perceptual reversal, while the same stimuli presented to the dominant eye decrease the reversal probability (Metzger et al., 2017). In our study, new fixation points were simultaneously presented to the dominant and suppressed eye. It is thus difficult to make a prediction regarding the modulation of perceptual states based on the existing literature. However, in the context of the generalized flash suppression paradigm, following adaptation to a monocular stimulus, the abrupt onset of the random dot motion in both eyes seems even more efficient to suppress the target than presenting them only to the non-adapted (and perceived) eye (Wilke et al., 2003). It is thus not unreasonable to assume that the binocular onset of new stimuli (Task 3) can induce a reversal too. Indeed, we found that the onset of new fixation points in both eyes can induce a reversal, however, the effect was considerably smaller than the one of FP shifts and saccade (Task 1 and 4). Thus, the mere appearance of new FPs on the rivaling objects has a milder effect on PSLs, being statistically significant in some analysis methods and not in others. In future studies with the current paradigm, the onsets of the fixation points could be made less salient by gradually fading them in, rather than turning them on abruptly. It is also possible that percepts would be more stable when adding some stimulus conditions that were used in the original study such as applying rotation filters and using slightly smaller stimuli (Hesse & Tsao, 2020).

### Using the novel no-report paradigm to study neural correlates of the perceptual switches

The original paradigm we employed in this experiment (Hesse & Tsao, 2020), is valuable because it is a no-report paradigm, where we can infer the perceived and suppressed object from the gaze locations of the subjects and use it to associate neural activity with either perceptual state. Although the Hesse and Tsao’s paradigm is well suited in terms of studying the neural differences between two perceived stimulus contents, switches between these two percepts do not necessarily occur spontaneously in this paradigm. Rather, saccades to the new FPs can induce a switch of the perceived stimulus. In our data, we found that around 14% of such saccades induce a perceptual switch within the expected latency range for a saccade-induced perceptual switch in the original paradigm. We also found that around 24% of all post-saccadic perceptual switches are “caused” by a preceding saccade. Given that conscious perception research is also trying to understand the neural underpinnings of spontaneous switches in perceptual states (Dwarakanath et al., 2023; Frässle et al., 2014; Kapoor et al., 2022), it is important that such a research question should be answered by using perceptual switches that occur *spontaneously*, rather than being induced by an external event.

We still believe that this no-report paradigm (Task 1) can be used for studying the mechanism and temporal dynamics of switches in conscious perception. One modification of the paradigm toward that end could be to increase the trial durations. Frequency of FP shifts can be decreased accordingly, lowering the number of saccades. Relatedly, future studies using the original paradigm from Hesse and Tsao (2020) to investigate the neural mechanisms behind spontaneous perceptual switches should perform pilot experiments, investigating the time limits during which subjects (monkeys or humans) can keep their gaze at the fixation point of a perceived stimulus, before performing a non-specific saccade due to fatigue, boredom and decrease in attention. In addition, events at a specific latency range around the spontaneous or task-induced saccades should be detected and potentially discarded from the neural correlates of spontaneous perceptual switches. An additional solution to investigate the neural mechanisms behind spontaneous alterations in perceptual states would be to only use secondary perceptual switches. In the current study, the total number of trials had to be kept low in Task 1 due to the presence of three more tasks. Still, subjects experienced a median of 22 secondary perceptual switches out of 72 complete perceptual switches, which means that approximately 30% of all complete switches were secondary events, and thus usable in the analyses of spontaneous switches (see **Table 3**). Another alternative to choosing the putative spontaneous perceptual switches would be to determine the time window after the trial onset in which perceptual switch probability is higher in the original paradigm (Task 1) compared to our control paradigm (Task 2). In **Figure 6B**, it can be seen that during the first 1.5 seconds following the trial onset, the perceptual switch probability is larger than in Task 2; which means that saccade requirements and location changes in the FPs could have an influence on this difference. One could exclude this time window, and use the perceptual switches that occur later than 1.5 seconds after the trial onset. In our dataset, the median ratio of perceptual switches that occurred after this time window is 63.5%, and the ratio increases to 64.5% if we categorize switches as spontaneous if they occur late (>1.5 seconds after trial onsets) *or* if they are secondary switches. One can also exclude trials based on the saccade onsets rather than the trial onsets, in order to decide on the subset of perceptual switches that are likely not induced by a preceding saccade. In our analysis with saccade-locked perceptual switches across all subjects, we found that perceptual switch probability is increased in the first 0.7 seconds following a saccade onset (see **Fig. 7B**). One could take all trials with a perceptual switch that is at least more than 0.7 seconds later than a saccade (an estimated 74.5% trials in our Task 1, see **Supplementary Table 3**), although these estimates are likely to be different in different subjects.

In summary, our data suggest that the original no-report paradigm from Hesse and Tsao (2020) should be treated carefully when the neural correlates of perceptual reversals are at the center of the study. Shifts in fixation points and particularly saccades often induce a perceptual reversal, but spontaneous perceptual switches can still be separated from these external factors by prolonging the periods between the shifts in fixation points of the objects, or excluding perceptual switch events around the onset of these trigger events. The results of our study point to the importance of using eye tracking for any study trying to elucidate the nature of endogenous switches during binocular rivalry, as any saccades, whether task-induced or involuntary, may exogenously increase the probability of perceptual switches.

### Authors contributions

Conceptualization: RS, IK, MW. Data curation: RS, IK. Data acquisition: RS. Stimuli and experimental advice: JH. Software: EK, RS, IK. Formal analysis: EK, RS. Funding acquisition: IK, MW. Project Administration: IK, MW. Supervision: IK, MW. Visualization: EK, IK. Writing—Original draft preparation: EK, IK, MW. Writing—Review & editing: EK, RS, JH, IK, MW.

## Acknowledgements

We thank Lukas Schneider, Daniela Lazzarini and Maryia Khvatava for technical and experimental support. We thank Doris Tsao for valuable discussions and for providing their stimulus material.

## Funding

This work was supported by the Marie Skłodowska-Curie Innovative Training Network: 2020-MSCA-ITN-2020 “In2PrimateBrains” (to MW and IK) and the German Research Foundation GRK2824 “Heart and Brain Diseases” (to MW and EK). The work was also supported by core funding of the Department of Cognitive Neurology (University Medical Center Goettingen) and the Cognitive Neuroscience Laboratory (German Primate Center).

### Declaration of generative AI and AI-assisted technologies in the writing process

During the preparation of this work we used ChatGPT 4 in order to facilitate some aspects of data analysis implementations in MATLAB. We also used it for instructions on how to use

Affinity tools. After using this tool/service, we reviewed and edited the code and take full responsibility for the content of the published article.

### Data and code availability statement

The datasets generated and analyzed for the current study, and the corresponding code, are available from the corresponding author on request. All code implementing the stimulus properties, the task, and the data analysis can be found in this public GitHub repository: https://github.com/dagdpz/BinoRiv_In2PB.

## References

Andersen, L. M., Vinding, M. C., Sandberg, K., & Overgaard, M. (2022). Task requirements affect the neural correlates of consciousness. The European Journal of Neuroscience, 56(10), 5810– 5822. 10.1111/ejn.15820

Aru, J., Bachmann, T., Singer, W., & Melloni, L. (2012). Distilling the neural correlates of consciousness. Neuroscience and Biobehavioral Reviews, 36(2), 737–746. 10.1016/j.neubiorev.2011.12.003

Bachmann, T., & Aru, J. (2023). Conscious interpretation: A distinct aspect for the neural markers of the contents of consciousness. Consciousness and Cognition, 108, 103471. 10.1016/j.concog.2023.103471

Blake, R., Fox, R., & McIntyre, C. (1971). Stochastic properties of stabilized-image binocular rivalry alternations. Journal of Experimental Psychology, 88(3), 327–332. 10.1037/h0030877

Blake, R., Westendorf, D., & Fox, R. (1990). Temporal perturbations of binocular rivalry. Perception & Psychophysics, 48(6), 593–602. 10.3758/bf03211605

Brainard, D. H. (1997). The Psychophysics Toolbox. Spatial Vision, 10(4), 433–436.

Brascamp, J., Sterzer, P., Blake, R., & Knapen, T. (2018). Multistable Perception and the Role of the Frontoparietal Cortex in Perceptual Inference. Annual Review of Psychology, 69, 77–103. 10.1146/annurev-psych-010417-085944

Carmel, D., Arcaro, M., Kastner, S., & Hasson, U. (2010). How to Create and Use Binocular Rivalry. Journal of Visualized Experiments, 45, 2030. 10.3791/2030

Carter, O., van Swinderen, B., Leopold, D. A., Collin, S. P., & Maier, A. (2020). Perceptual rivalry across animal species. The Journal of Comparative Neurology, 528(17), 3123–3133. 10.1002/cne.24939

Dehaene, S., & Changeux, J.-P. (2011). Experimental and Theoretical Approaches to Conscious Processing. Neuron, 70(2), 200–227. 10.1016/j.neuron.2011.03.018

Duman, I., Ehmann, I. S., Gonsalves, A. R., Gültekin, Z., Van den Berckt, J., & van Leeuwen, C. (2022). The No-Report Paradigm: A Revolution in Consciousness Research? Frontiers in Human Neuroscience, 16. 10.3389/fnhum.2022.861517

Dwarakanath, A., Kapoor, V., Werner, J., Safavi, S., Fedorov, L. A., Logothetis, N. K., & Panagiotaropoulos, T. I. (2023). Bistability of prefrontal states gates access to consciousness. Neuron, 111(10), 1666–1683.e4. 10.1016/j.neuron.2023.02.027

Fox, R., & Check, R. (1968). Detection of motion during binocular rivalry suppression. Journal of Experimental Psychology, 78(3), 388–395. 10.1037/h0026440

Frässle, S., Sommer, J., Jansen, A., Naber, M., & Einhäuser, W. (2014). Binocular rivalry: Frontal activity relates to introspection and action but not to perception. The Journal of Neuroscience: The Official Journal of the Society for Neuroscience, 34(5), 1738–1747. 10.1523/JNEUROSCI.4403-13.2014

Gail, A., Brinksmeyer, H. J., & Eckhorn, R. (2004). Perception-related modulations of local field potential power and coherence in primary visual cortex of awake monkey during binocular rivalry. *Cerebral Cortex (New York*, N.Y.: 1991), *14*(3), 300–313. 10.1093/cercor/bhg129

Gelbard-Sagiv, H., Mudrik, L., Hill, M. R., Koch, C., & Fried, I. (2018). Human single neuron activity precedes emergence of conscious perception. Nature Communications, 9(1), 2057. 10.1038/s41467-018-03749-0

Hesse, J. K., & Tsao, D. Y. (2020). A new no-report paradigm reveals that face cells encode both consciously perceived and suppressed stimuli. eLife, 9, e58360. 10.7554/eLife.58360

Joshi, S., & Gold, J. I. (2020). Pupil size as a window on neural substrates of cognition. Trends in Cognitive Sciences, 24(6), 466–480. 10.1016/j.tics.2020.03.005

Kapoor, V., Dwarakanath, A., Safavi, S., Werner, J., Besserve, M., Panagiotaropoulos, T. I., & Logothetis, N. K. (2022). Decoding internally generated transitions of conscious contents in the prefrontal cortex without subjective reports. Nature Communications, 13(1), 1535. 10.1038/s41467-022-28897-2

Keliris, G. A., Logothetis, N. K., & Tolias, A. S. (2010). The role of the primary visual cortex in perceptual suppression of salient visual stimuli. The Journal of Neuroscience: The Official Journal of the Society for Neuroscience, 30(37), 12353–12365. 10.1523/JNEUROSCI.0677-10.2010

Kloosterman, N. A., Meindertsma, T., van Loon, A. M., Lamme, V. A. F., Bonneh, Y. S., & Donner, T. H. (2015). Pupil size tracks perceptual content and surprise. European Journal of Neuroscience, 41(8), 1068–1078. 10.1111/ejn.12859

Kornmeier, J., & Bach, M. (2012). Ambiguous Figures – What Happens in the Brain When Perception Changes But Not the Stimulus. Frontiers in Human Neuroscience, 6. https://www.frontiersin.org/articles/10.3389/fnhum.2012.00051

Lempert, K. M., Chen, Y. L., & Fleming, S. M. (2015). Relating Pupil Dilation and Metacognitive Confidence during Auditory Decision-Making. PLOS ONE, 10(5), e0126588. 10.1371/journal.pone.0126588

Leopold, D. A., & Logothetis, N. K. (1996). Activity changes in early visual cortex reflect monkeys’ percepts during binocular rivalry. Nature, 379(6565), 549–553. 10.1038/379549a0

Leopold, D. A., & Logothetis, N. K. (1999). Multistable phenomena: Changing views in perception. Trends in Cognitive Sciences, 3(7), 254–264. 10.1016/s1364-6613(99)01332-7

Leopold, D. A., Plettenberg, H. K., & Logothetis, N. K. (2002). Visual processing in the ketamine-anesthetized monkey. Optokinetic and blood oxygenation level-dependent responses. Experimental Brain Research, 143(3), 359–372. 10.1007/s00221-001-0998-0

Lepauvre, A., & Melloni, L. (2021). The search for the neural correlate of consciousness: Progress and challenges. Philosophy and the Mind Sciences, 2. 10.33735/phimisci.2021.87

Logothetis, N. K., & Schall, J. D. (1989). Neuronal correlates of subjective visual perception. *Science (New York*, N.Y.), 245(4919), 761–763. 10.1126/science.2772635

Lumer, E. D., Friston, K. J., & Rees, G. (1998). Neural correlates of perceptual rivalry in the human brain. *Science (New York*, N.Y.), 280(5371), 1930–1934. 10.1126/science.280.5371.1930

Maier, A., Logothetis, N. K., & Leopold, D. A. (2007). Context-dependent perceptual modulation of single neurons in primate visual cortex. Proceedings of the National Academy of Sciences of the United States of America, 104(13), 5620–5625. 10.1073/pnas.0608489104

Metzger, B. A., & Beck, D. M. (2020). Probing the mechanisms of probe-mediated binocular rivalry. Vision Research, 173, 21–28. 10.1016/j.visres.2020.04.011

Metzger, B. A., Mathewson, K. E., Tapia, E., Fabiani, M., Gratton, G., & Beck, D. M. (2017). Regulating the Access to Awareness: Brain Activity Related to Probe-related and Spontaneous Reversals in Binocular Rivalry. Journal of Cognitive Neuroscience, 29(6), 1089– 1102. 10.1162/jocn_a_01104

Naber, M., Frässle, S., & Einhäuser, W. (2011). Perceptual rivalry: Reflexes reveal the gradual nature of visual awareness. PloS One, 6(6), e20910. 10.1371/journal.pone.0020910

Panagiotaropoulos, T. I., Deco, G., Kapoor, V., & Logothetis, N. K. (2012). Neuronal Discharges and Gamma Oscillations Explicitly Reflect Visual Consciousness in the Lateral Prefrontal Cortex. Neuron, 74(5), 924–935. 10.1016/j.neuron.2012.04.013

Pettigrew, J. D. (2001). Searching for the Switch: Neural Bases for Perceptual Rivalry Alternations. Brain and Mind, 2(1), 85–118. 10.1023/A:1017929617197

Ross, J., & Ma-Wyatt, A. (2004). Saccades actively maintain perceptual continuity. Nature Neuroscience, 7(1), 65–69. 10.1038/nn1163

Sheinberg, D. L., & Logothetis, N. K. (1997). The role of temporal cortical areas in perceptual organization. Proc. Natl. Acad. Sci. USA.

Storm, J. F., Boly, M., Casali, A. G., Massimini, M., Olcese, U., Pennartz, C. M. A., & Wilke, M. (2017). Consciousness Regained: Disentangling Mechanisms, Brain Systems, and Behavioral Responses. The Journal of Neuroscience, 37(45), 10882–10893. 10.1523/JNEUROSCI.1838-17.2017

Storm, J. F., Klink, P. C., Aru, J., Senn, W., Goebel, R., Pigorini, A., Avanzini, P., Vanduffel, W., Roelfsema, P. R., Massimini, M., Larkum, M. E., & Pennartz, C. M. A. (2024). An integrative, multiscale view on neural theories of consciousness. Neuron, 112(10), 1531–1552. 10.1016/j.neuron.2024.02.004

Tong, F., Meng, M., & Blake, R. (2006). Neural bases of binocular rivalry. Trends in Cognitive Sciences, 10(11), 502–511. 10.1016/j.tics.2006.09.003

Tsuchiya, N., Frässle, S., Wilke, M., & Lamme, V. (2016). No-Report and Report-Based Paradigms Jointly Unravel the NCC: Response to Overgaard and Fazekas. Trends in Cognitive Sciences, 20(4), 242–243. 10.1016/j.tics.2016.01.006

Tsuchiya, N., Wilke, M., Frässle, S., & Lamme, V. A. F. (2015). No-Report Paradigms: Extracting the True Neural Correlates of Consciousness. Trends in Cognitive Sciences, 19(12), 757–770. 10.1016/j.tics.2015.10.002

Urai, A. E., Braun, A., & Donner, T. H. (2017). Pupil-linked arousal is driven by decision uncertainty and alters serial choice bias. Nature Communications, 8(1), 14637. 10.1038/ncomms14637

Van Dam, L. C. J., & Van Ee, R. (2006a). Retinal image shifts, but not eye movements per se, cause alternations in awareness during binocular rivalry. Journal of Vision, 6(11), 3–3. 10.1167/6.11.3

Van Dam, L. C. J., & Van Ee, R. (2006b). The role of saccades in exerting voluntary control in perceptual and binocular rivalry. Vision Research, 46(6–7), 787–799. 10.1016/j.visres.2005.10.011

Wade, N. J. (1974). The effect of orientation in binocular contour rivalry of real images and afterimages. Perception & Psychophysics, 15(2), 227–232. 10.3758/BF03213937

Wilke, M., Logothetis, N. K., & Leopold, D. A. (2003). Generalized Flash Suppression of Salient Visual Targets. Neuron, 39(6), 1043–1052. 10.1016/j.neuron.2003.08.003

Wilke, M., Mueller, K.-M., & Leopold, D. A. (2009). Neural activity in the visual thalamus reflects perceptual suppression. Proceedings of the National Academy of Sciences, 106(23), 9465– 9470. 10.1073/pnas.0900714106

Zhang, R., Engel, S. A., & Kay, K. (2017). Binocular Rivalry: A Window into Cortical Competition and Suppression. Journal of the Indian Institute of Science, 97(4), 477–485. 10.1007/s41745-017-0048-y

