## Supplementary Information for "Fixation shifts in a novel “no-report” binocular rivalry paradigm induce saccade-related perceptual switches"

Supplementary information contains:

**Supplementary Results SR.1 – SR.4**

**10 Supplementary Figures**

**3 Supplementary Tables**

#### Supplementary Results

##### SR.1. Perceptual Switch Latency (PSL) analysis with only complete and only incomplete perceptual switches

Perceptual switches that the subjects reported were of two kinds: a switch in perception from one object to the other (complete perceptual switch), or a temporary interruption in the perception of one of the objects (incomplete perceptual switch, see **Figure 1** and **Methods** for a detailed description). To supplement our behavioral analyses, we repeated the PSL analyses (see **Figures 2, 3** and **4**) while selecting only the complete perceptual switches. We observed that the pooled PSL distributions in Task 1 and Task 4 differ significantly from the respective uniform distributions (two-sample Kolmogorov-Smirnov tests; $p=4.55x10^{-4}$ and $p=5.30x10^{-4}$, respectively), while Task 2 and Task 3 did not differ from uniform distribution (p=0.67 and p=0.04, respectively, **Supplementary Figure 3**, right column). Group-level distributions of median complete PSLs did not differ significantly from the theoretically expected 2.5 seconds (see **Supplementary Figure 2** for PSL simulations) in any of the tasks: Task 1 (p=0.06), Task 2 (p=0.72), Task 3 (p=0.04), and Task 4 (p=0.01) all larger than the corrected $\alpha$ = 0.0125. Note that in all tasks except Task 2, there is a trend towards median PSL values being less than 2.5, but these trends did not reach statistical significance when only the complete perceptual switches are used (**Supplementary Figure 3**, left column).

**Supplementary Figure 4** displays a similar analysis as in **Supplementary Figure 3**, while including only the incomplete perceptual switches. We performed the same statistical tests as we did for all (**Figure 3**) and only complete perceptual switches (**Supplementary Figure 3**), but the number of incomplete switches that the subjects reported were lower than the number of complete switches. Thus, **Supplementary Figure 4** serves more as a qualitative picture of how the latencies of incomplete perceptual switches (the “momentary” interruptions in between periods of perceiving the same object) look like. Of note is that in Tasks 1 and 4, the PSLs appear more time-locked to the trial onsets than in Tasks 2 and 3, suggesting that the saccade instruction/requirement at trial onsets do not just induce complete perceptual switches but also interrupt perceptual dominance periods specifically at the beginning of trials; which likely changes the time course of the perceptual dominance epochs in these BR tasks. We also compared the number of incomplete switches that each subject experienced across tasks (see raw values in **Supplementary Table 1**), and found that task identity has an overall effect on this number (one-way repeated measures Friedman's test; $\chi 2=9.53$, $p=0.02$). Post-hoc tests between task pairs revealed that the subjects experienced significantly more incomplete perceptual switches in Task 1 compared to Task 2 ([Mean SD]: Task 1 = [17.78 14.52], Task 2 = [13.24 13.06]; p=0.02), showing that the number of such 'interruptions' in perception is increased in the original BR paradigm (Task 1) with respect to our negative control experiment (Task 2). However, the ratio of incomplete switches to all perceptual switches (complete and incomplete) is not significantly different between tasks (one-way repeated measures Friedman's test; $\chi 2=5.21$, $p=0.16$), showing that both incomplete and complete perceptual switch events increased in a similar amount in Tasks 1 and 2, rather than a specific increase in the incomplete switch events.

#### SR.2. Perceptual dominance durations

**Supplementary Figure 5** shows the perceptual dominance durations (i.e., lengths of time windows where the subject reported perceiving only one object without interruptions or changes in perception) from two example subjects (**A-B**), and pooled across all subjects in each task (**C-F**). The average perceptual dominance durations of each subject in each task are also reported in **Supplementary Table 1**. It is apparent that there are numerous perceptual dominance durations that are smaller than or equal to 3-4 seconds. This means that whenever an external event (at the trial onset) induces a perceptual switch early on within a 5-second trial, a secondary perceptual switch might occur before the trial ends, simply because the perceptual dominance durations are not long enough. This would introduce PSL values later on in the trial, possibly accounting for the median PSLs getting closer to the mid-point of the trials (**Figure 3**, left column). Relatedly, we analyzed how the early perceptual switches affect further (secondary) perceptual switches within the same trials. We took the subset of all trials that included a perceptual switch within the first second and found that PSLs in the remaining trials (from 1 to 5 seconds since trial onset) show a significantly different distribution than a uniform distribution of the same size (two-sample Kolmogorov-Smirnov test, $p=1.36\times10^{-3}$), with a higher probability of switch events closer to the end of these trials (**Supplementary Figure 5 G-H**). Inevitably, we include these secondary events while calculating the median PSLs of each individual subject. This provides a potential explanation to why the distributions of individual median PSLs at the group-level show a relatively weak "time-locking effect" (**Figure 3, left column**), despite still being statistically significant after correction for multiple comparisons in Tasks 1 and 4 (**Figure 3A, B, G, H**).

**Supplementary Figure 6** displays the time spent in piecemeal perception for each subject in each task during the binocular rivalry condition, and compares these durations with the reaction times of the subjects to report a perceptual switch from one stimulus to the other in the physical control condition. Some subjects seemed to be experiencing piecemeal percepts for significant amounts of time (>1 second on average), whereas most subjects did not report piecemeal perception based on their similar reaction times of switching from one percept to the other in binocular rivalry and physical conditions.

#### SR.3. Latencies with peak probability for perceptual switches

As a strategy to isolate the effect of external factors (saccade and shifting FPs) on PSLs, we also analyzed the maximum latency probabilities of each individual in each task. The individual-level histograms are exemplified in **Supplementary Figure 7**. We chose a bin size of 0.25 seconds, determined the latency bin with the maximum probability, and used the mean time point of that bin as the peak latency value of that subject in the given task (e.g., if the latency bin with the maximum probability is 0.25-0.50 s, the peak latency value is 0.375 s). If there are multiple bins that have the same maximum value, the resulting peak value is determined as the average of these latency bins.

Peak latencies were compared across tasks at the group-level, in a similar analysis as shown for overall median latencies (**Figure 2B**) and median latencies of the first switches (**Figure 5B**). One-way Friedman's test across tasks showed that task identity had a main effect on the peak PSLs at the group-level ($\chi^2$=12.17, p=6.84x$10^{-3}$; **Supplementary Figure 8B**). Post-hoc tests revealed that task pairs 1-2 ([Mean SD]: Task 1 = [1.16 0.94], Task 2 = [2.57 1.33]; p=4.92x$10^{-3}$, 95% CI = [-2.56 -0.33]) and 2-4 ([Mean SD]: Task 4 = [1.48 1.33]; p=0.05 (0.048 after Tukey-Cramer correction to be precise), 95% CI = [0.00 2.23]) had significantly different PSLs, showing that the peak PSLs were significantly longer in Task 2 than any other task. Grouping the tasks according to the presence of saccade requirements and shifting fixation points showed that the saccade requirement has a main effect on peak PSLs (**Supplementary Figure 8C**; $\chi^2$=7.33, p=6.80x$10^{-3}$), whereas shifting FPs does not (**Supplementary Figure 8D**; $\chi^2$=2.54, p=0.11).

In the absence of saccades or shifting FPs, the expected peak latency at the group-level is again 2.5 seconds. Thus, peak latencies from each task were separately compared against the 2.5 seconds by two-tailed Wilcoxon signed-rank tests (**Supplementary Figure 9A-D**). Peak PSLs at the group-level were significantly closer to the trial onset than 2.5 seconds in Task 1 ([Mean SD] = [1.16 0.94]; left-tailed Wilcoxon signed-rank test against 2.5 seconds; p=1.13x$10^{-3}$), but not in Task 2 ([Mean SD] = [2.57 1.33]; p=0.83) or Task 3 ([Mean SD] = [1.77 1.28]; p=0.03, trend does not reach statistical significance), and Task 4 ([Mean SD] = [1.48 1.33]; p=0.01, trend does not reach statistical significance (p>0.0125)). When the peak PSLs in each task are compared to Task 2 (negative control) with paired Wilcoxon signed-rank tests, Tasks 1 and 4 have a significantly smaller peak PSLs than that of Task 2 (p=2.85x$10^{-3}$, and p=7.40x$10^{-3}$, respectively) while Task 3 did not have peak latencies significantly different from Task 2 (p=0.08; **Supplementary Figure 9E-G**). Distributions of peak PSLs, specifically in Tasks 1 and 4, display a qualitatively stronger time-locking to the trial onsets when compared to the weaker effect that we observed by investigating the median PSLs (**Figure 3**, left).

#### SR.4. Perceptual switches in [0 1.5] s of trials with fixation point shifts (perceptual switch rates putatively caused by the external factors at trial onsets)

We expanded our analyses based on the estimation that perceptual switches within the first 1.5 seconds following the trial onset are likely to be induced by the external factors that take place at the moment of the trial onset (saccade instruction and/or shifting FPs) (see **Figure 6B**). We first analyzed the percentage of trials with a perceptual switch within the first 1.5 seconds after the location shift of the fixation points. Comparing this percentage across tasks revealed that task identity had a main effect on the percentage of trials with a switch within the first 1.5 seconds (one-way Friedman's test; $\chi^2$= 18.25, p=3.91x$10^{-4}$). Post-hoc tests with correction for multiple comparisons revealed that task pairs 2-4 ([Mean SD]: Task 4 = [62.46 18.42], Task 2 = [38.01 12.66]; p=6.76x$10^{-4}$, 95% CI = [-2.84 -0.57]) and 3-4 ([Mean SD]: Task 3 = [42.08 15.26]; p=3.10x$10^{-3}$, 95% CI = [-2.67 -0.39]) differed significantly (**Supplementary Figure 10A**). Even though the post-hoc comparison between Task 1 and Task 2 did not yield a statistically significant difference (p=0.25), 50.41% of trials in Task 1 had a perceptual switch within the first 1.5 seconds; which is 12.40% higher than in Task 2. This difference could be interpreted as the additional rate of trials where a perceptual switch is induced due to the external factors at the trial onsets. Finally, we compared the percentage of all perceptual switches within the first 1.5 seconds. We found that the task identity has a main effect in the ratio of perceptual switches within the first 1.5 seconds (one-way Friedman's test; $\chi^2$=27.28, p=5.14x$10^{-6}$). Post-hoc tests revealed that task pairs 1-2 ([Mean SD]: Task 1 = [62.27 14.03], Task 2 = [45.86 8.61]; p=0.02, 95% CI = [0.16 2.43]), 2-4 ([Mean SD]: Task 4 = [69.13 14.74]; p=1.98x$10^{-5}$, 95% CI = [-3.20 -0.92]), and 3-4 ([Mean SD]: Task 3 = [49.24 12.32]; p=3.92x$10^{-4}$, 95% CI = [-2.90 -0.63]) had significantly different ratios of early perceptual switches to all perceptual switches (**Supplementary Figure 10B**). In particular, the 16.41% increase in the ratio of early perceptual switches in Task 1 compared to Task 2 further shows that perceptual switch onsets are affected by external factors at the trial onsets.

Trials with fixation point shifts are also used for saccade detection at trial onsets, since the subject needed to find the new fixation point of the perceived object. **Supplementary Table 3** shows the number of trials with fixation point shifts at trial onsets, the number of detected saccades, and the number of estimated percentage of saccades that "cause" a perceptual switch at an early time window (see **Section 3.4** and **Figure 7**).

### Supplementary Figures

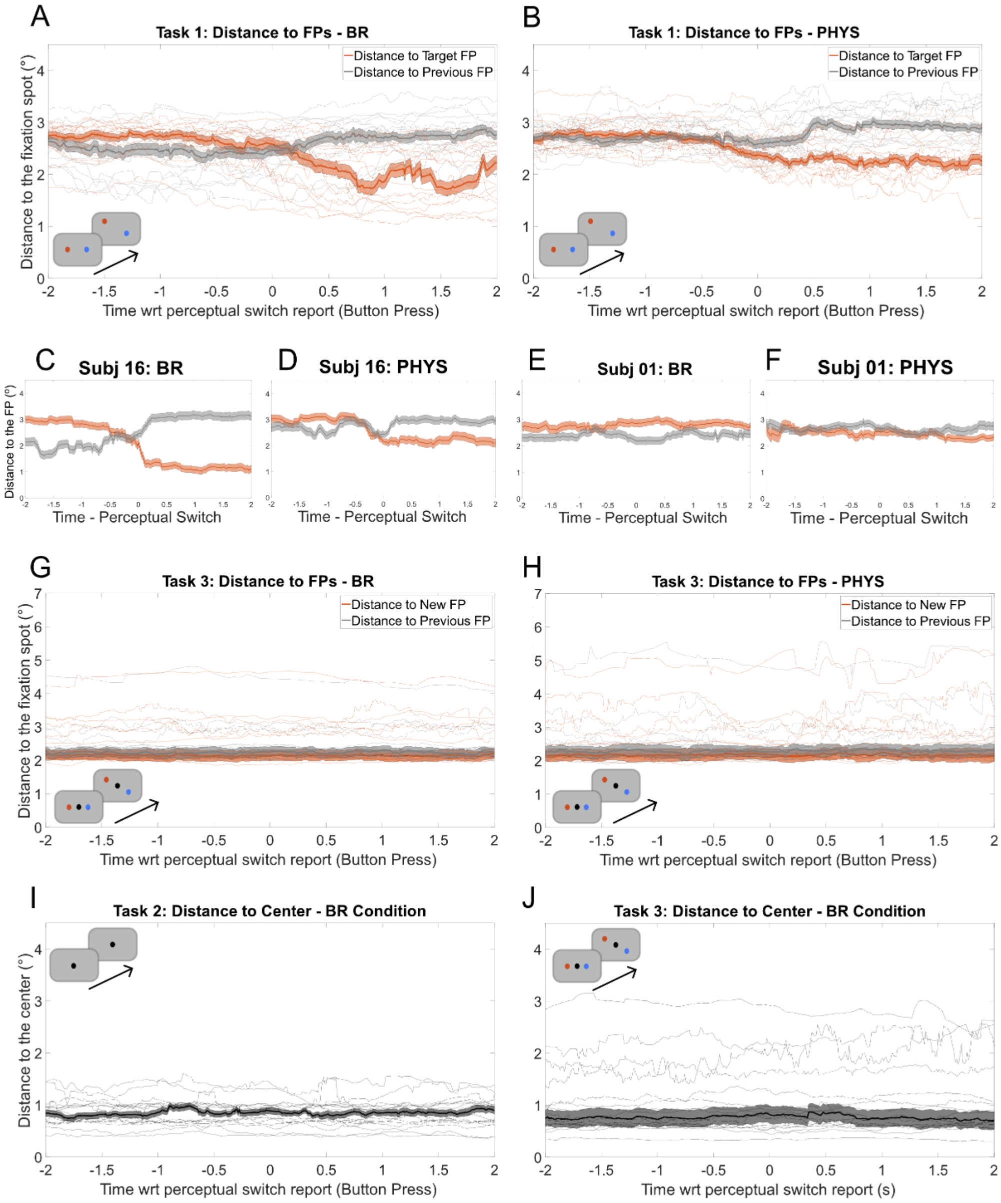


**Supplementary Figure 1: Eye-tracking analysis. (A-B)** Qualitative visualization of gaze distance to target and non-target fixation points (FP) in Task 1. Target FP denotes the FP corresponding to the object that the subject reported perceiving from time=0 onwards. In Task 1, the gaze locations are closer to the target FP after time point zero at the group level (as expected), both in the binocular rivalry **(A)** and physical stimulus **(B)** conditions. While some subjects yielded relatively accurate gaze tracking data **(C, D)**, it was not possible to infer the perceived object from the gaze data of each subject **(E, F)**. In Task 3, subjects were asked to fixate at the center of the screen and not at the jumping FPs. We show that the gaze locations do not get closer to the new FP (the FP corresponding to the object that is perceived from time point zero and onwards) in Task 3, which is the expected behavior in this task (**G** and **H**; note the contrast with panels **A-B**, which show that the gaze distance gets closer to the target FP and farther from the previous FP in Task 1). In Tasks 2 and 3, where the subjects always needed to fixate at the center of the screen, we checked that the gaze distance to the center does not change significantly at the moments of perceptual switches (**I** and **J**). In panels **A, B, G, H, I,** and **J**; each thin curve corresponds to the average data from a single subject.

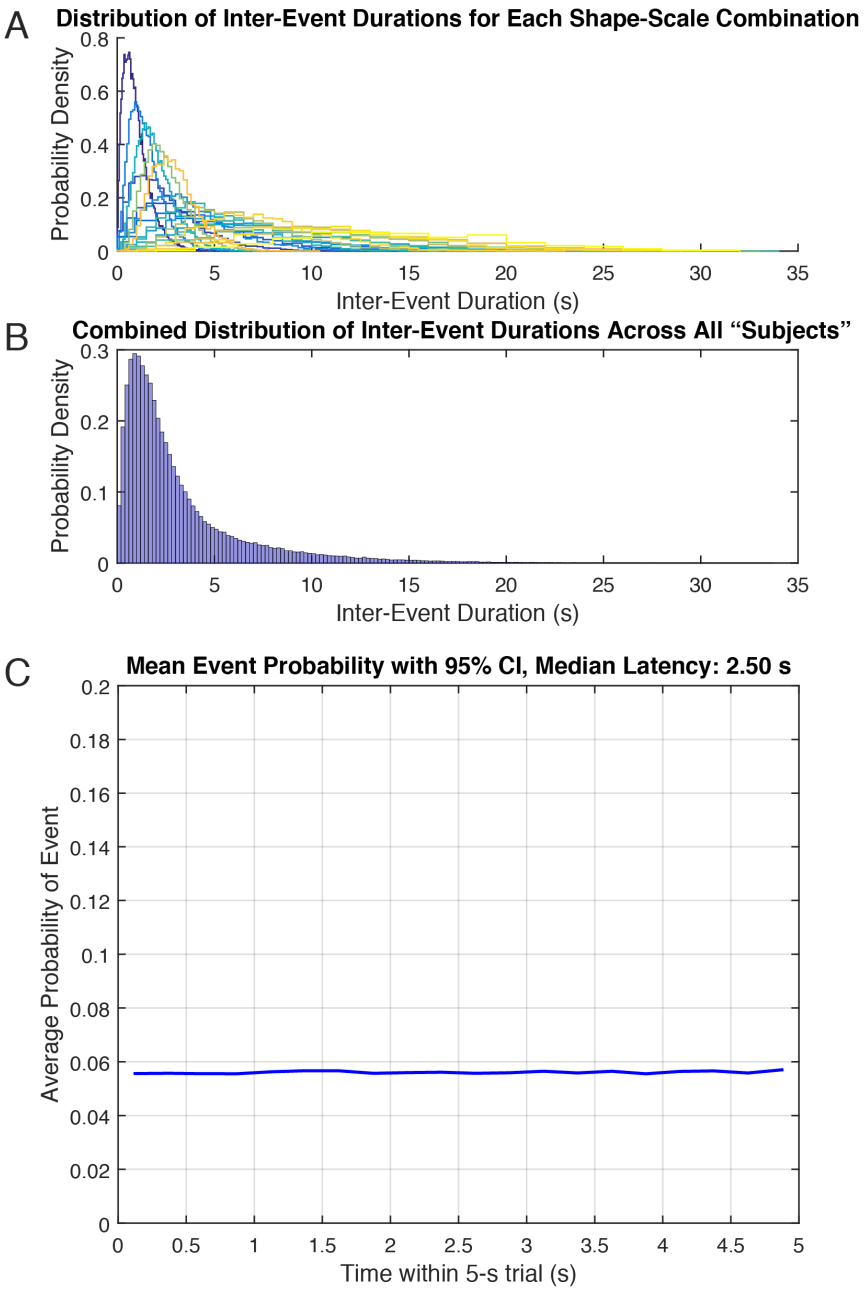


**Supplementary Figure 2: Perceptual Switch Latency (PSL) simulations.** (**A**) To simulate the PSL distributions within 5-second trials across subjects with different perceptual dominance duration distribution, we approximated each “subject’s” perceptual dominance duration distribution (inter-event durations) with a gamma distribution with a shape (ranging from 2 to 6) and scale (ranging from 0.5 to 3) parameters. (**B**) Group-level perceptual dominance duration histogram (after pooling all “subjects”) is also a gamma distribution. (**C**) When events are placed on the time axis of imaginary experimental blocks of continuous 5-second trials (as in our experimental Task 2), the probability distributions of perceptual switch latencies follow a uniform distribution throughout a 5-second trial, with a median latency of 2.50 seconds. This is the basis for our expectation that the experimental median PSL values of subjects should not be different from 2.50 seconds, in the absence of any external triggers.

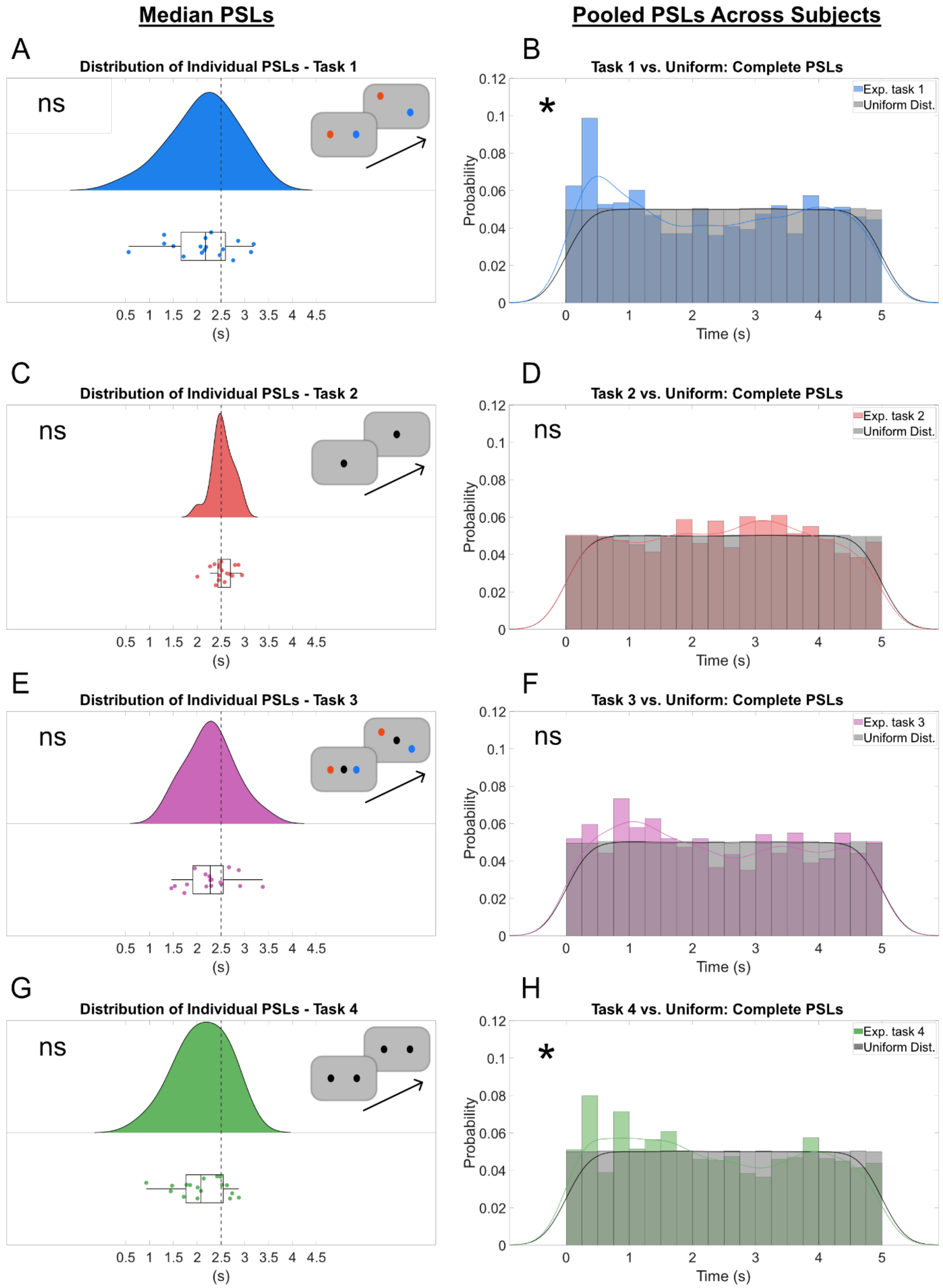


**Supplementary Figure 3: Single-task median PSL comparison against 2.5 seconds (left column) and comparison of PSLs pooled across all subjects against a uniform distribution (right column).** The analyses are identical to those shown in Figure 3, but here we use only the ***complete*** perceptual switches. A trend towards a time-locking effect in tasks 1,3, and 4 does not reach statistical significance, whereas pooling PSLs across subjects reveals a time-locking effect in Tasks 1 and 4, whereas the distributions do not differ from the expected null distributions in Tasks 2 and 3.

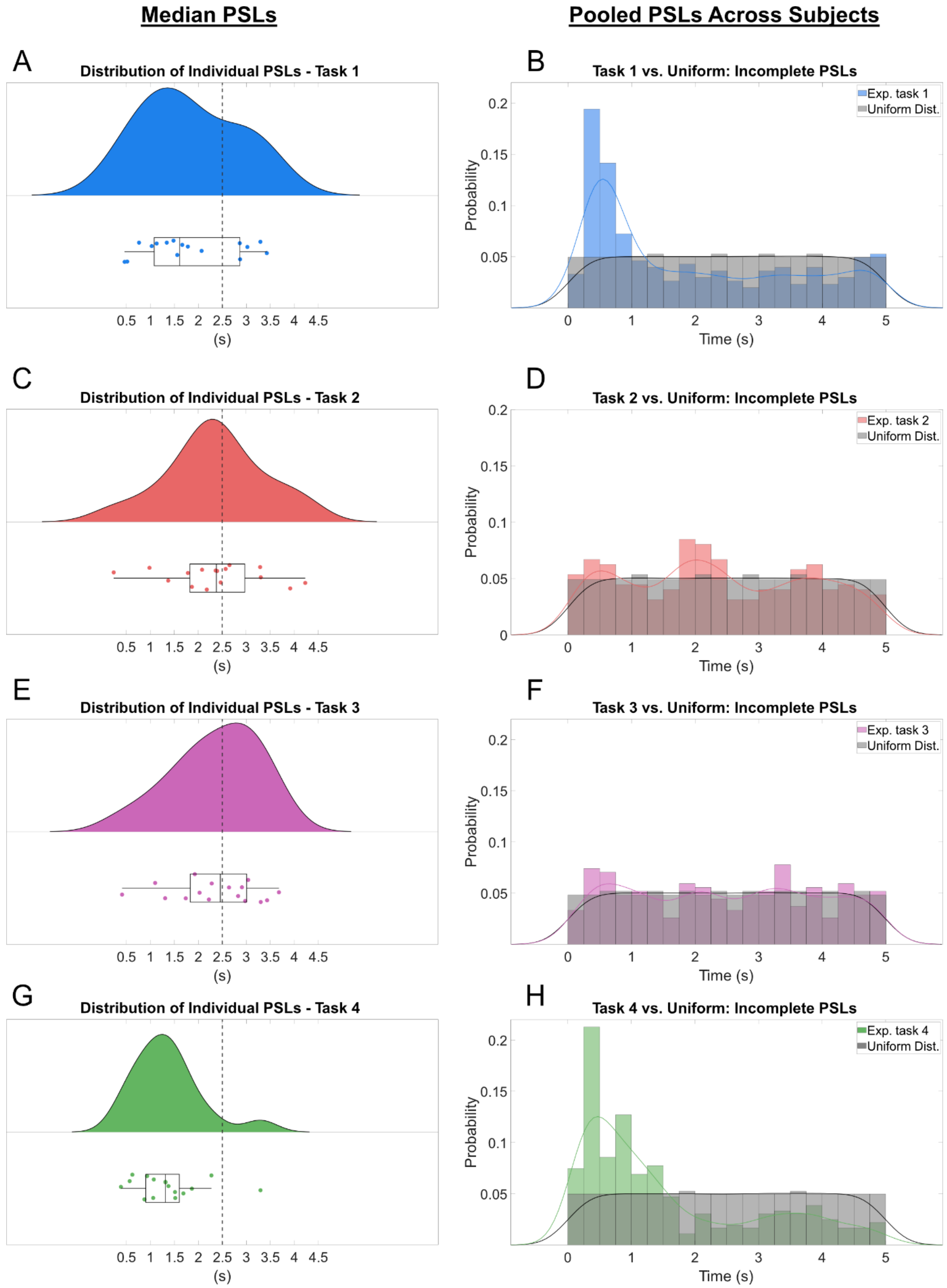


**Supplementary Figure 4: Single-task median PSL comparison against 2.5 seconds (left column) and comparison of PSLs pooled across all subjects against a uniform distribution (right column).** The analyses are identical to those shown in **Figure 3**, but here we use only the ***incomplete*** perceptual switches. Both types of latency distributions show a time-locking effect in Tasks 1 and 4, whereas the distributions do not differ from the expected null distributions in Tasks 2 and 3.

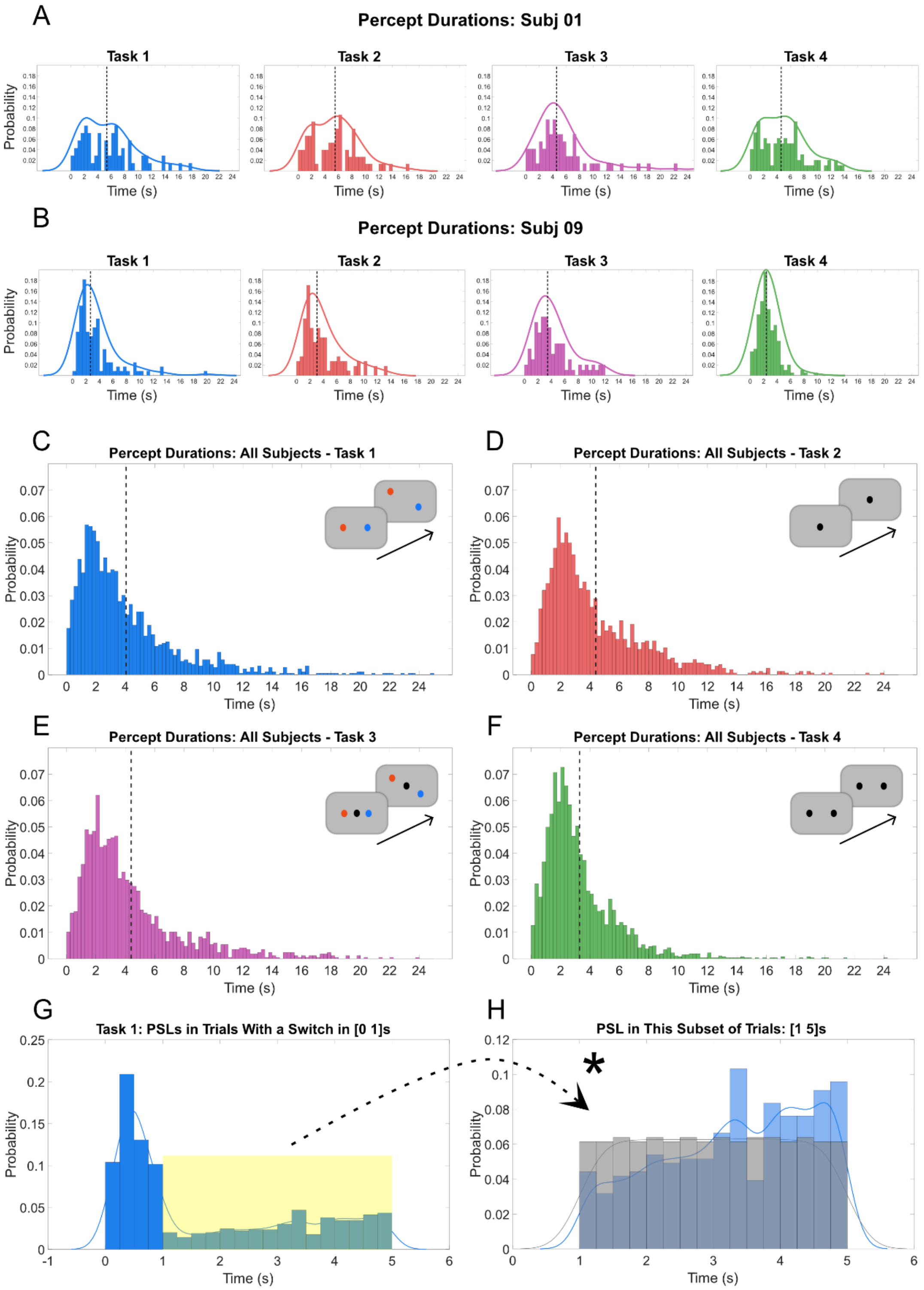


**Supplementary Figure 5: Perceptual dominance durations. (A-B)** Distribution of perceptual dominance durations from two example subjects. **(C-F)** Perceptual dominance durations pooled across all subjects in each task. Dashed vertical lines indicate the overall mean value of all dominance durations on the histograms. With the idea that trial-onset-induced perceptual switches (putatively the subset of perceptual switches that occur within the first second of a trial) might introduce a higher likelihood of secondary perceptual switches towards the end of trial, we took the subset of trials with a switch within the first second **(G)**, and compared the PSL distribution in the time window of 1 to 5 seconds to a uniform distribution of the same size **(H)**. The PSL distribution here is found to be significantly different from the uniform distribution, with a higher likelihood of perceptual switches towards the end of these trials.

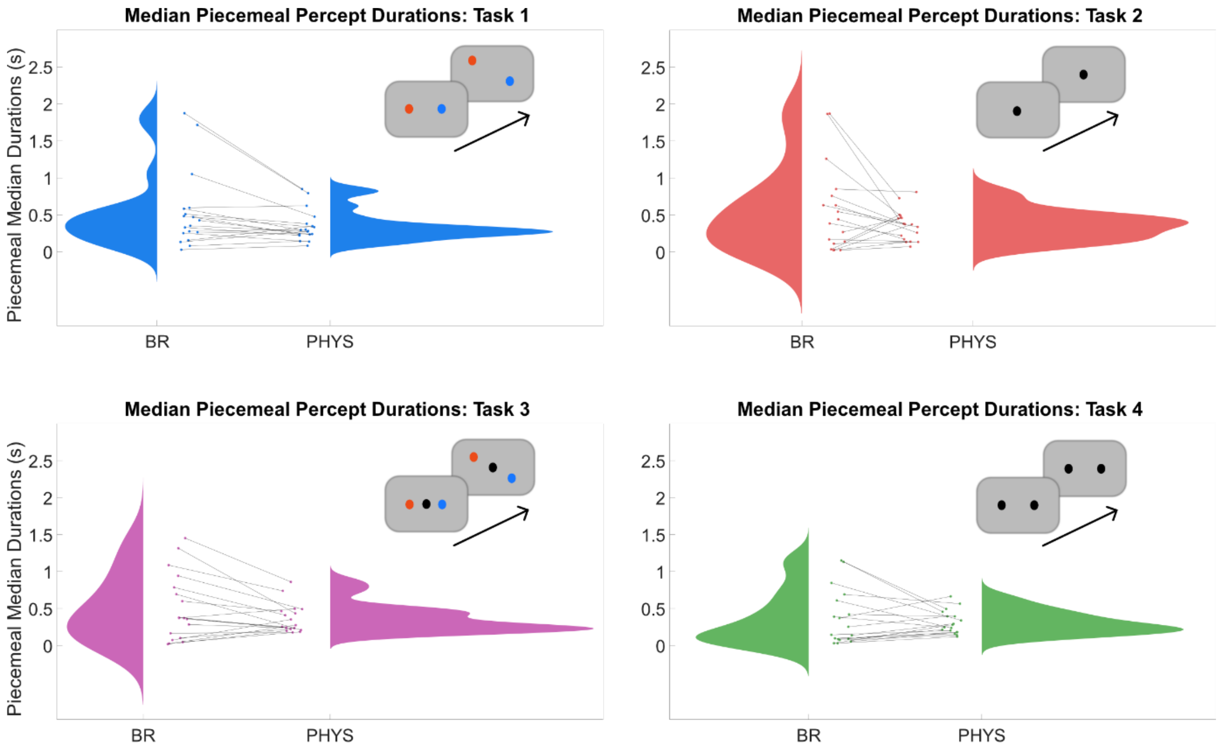


**Supplementary Figure 6: Piecemeal percept durations.** We aimed to calculate the time spent in the state of "piecemeal" percept (perception of both objects at the same time) during binocular rivalry, namely the durations when subjects pressed both buttons or none (see **Figure 1**). The median durations of such periods are obtained from each subject to create a group-level distribution of piecemeal percept durations. We did not use mixed pictures in the physical control condition, thus the "piecemeal" reports in the physical condition represent the reaction time required to report perceptual switches. By subtracting these times in the physical condition from those in the BR condition, we aimed to infer the actual time spent in piecemeal perception during binocular rivalry. We see from the differences between BR and the physical conditions in each task that some subjects perceived both objects ("piecemeal") for significant durations (piecemeal durations are much higher in the BR condition), whereas other subjects report direct switches from one object to the other without piecemeal perceptions in between (very similar "piecemeal" durations in BR and physical conditions).

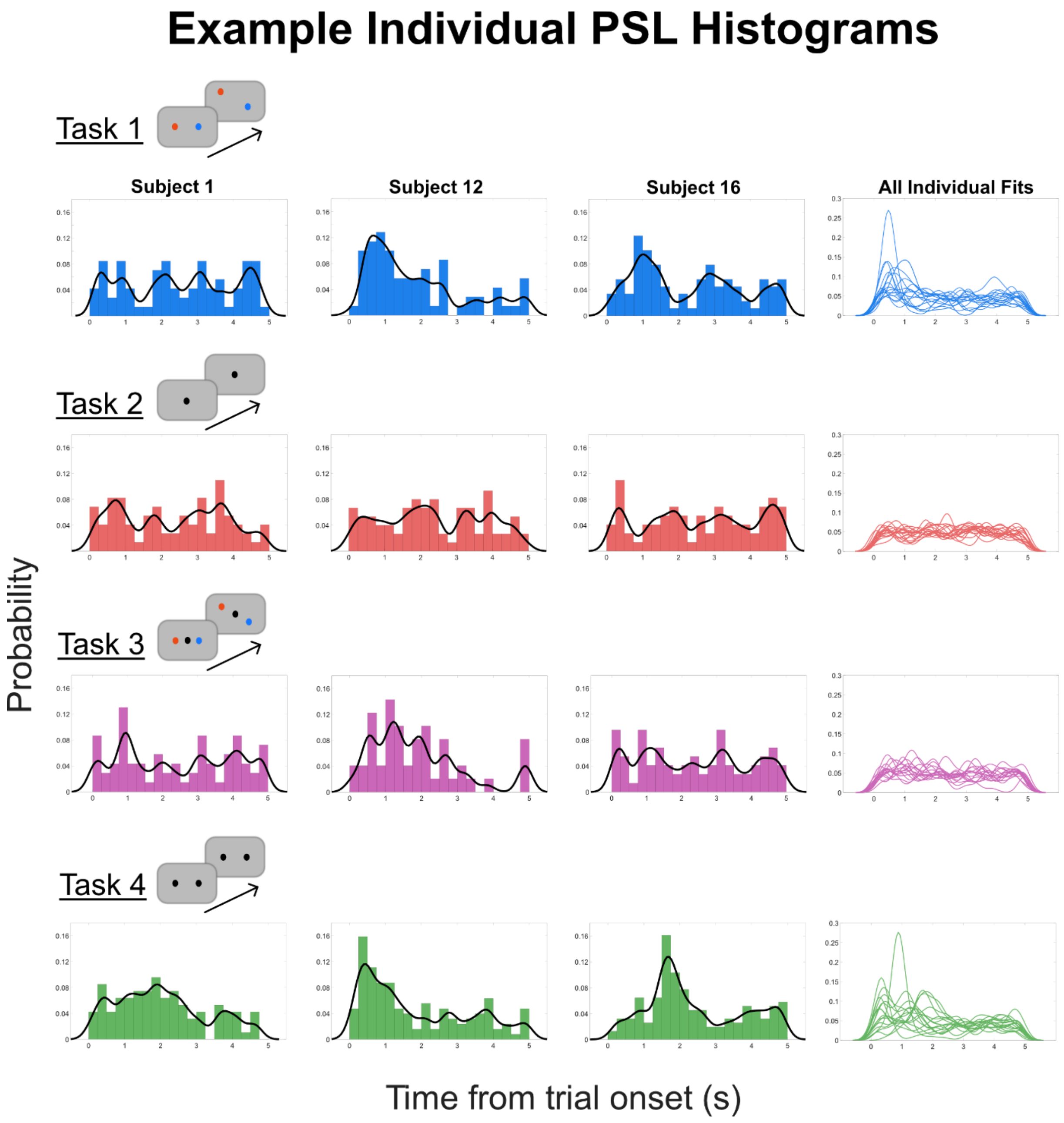


**Supplementary Figure 7: Subject-level PSL distributions.** Individual Perceptual Switch Latency (PSL) histograms from example individuals (subjects 1, 12, and 16). These subjects were chosen as qualitative examples representing different types of time-locking profiles to event onsets: subject 1 shows a relatively uniform probability distributions over the course of trials in each task (no time-locking), subject 12 shows time-locking effects in each task but task 2 (our "negative control" task), and subject 16 shows time-locking effects in Tasks 1 and 4, while not showing a prominent time-locking in Tasks 2 and 3; possibly contributing to the finding of only marginal time-locking effect of Task 3 at the group-level. On the right-most column, we used kernel density estimations for each individual PSL distribution from the 17 subjects in each task.

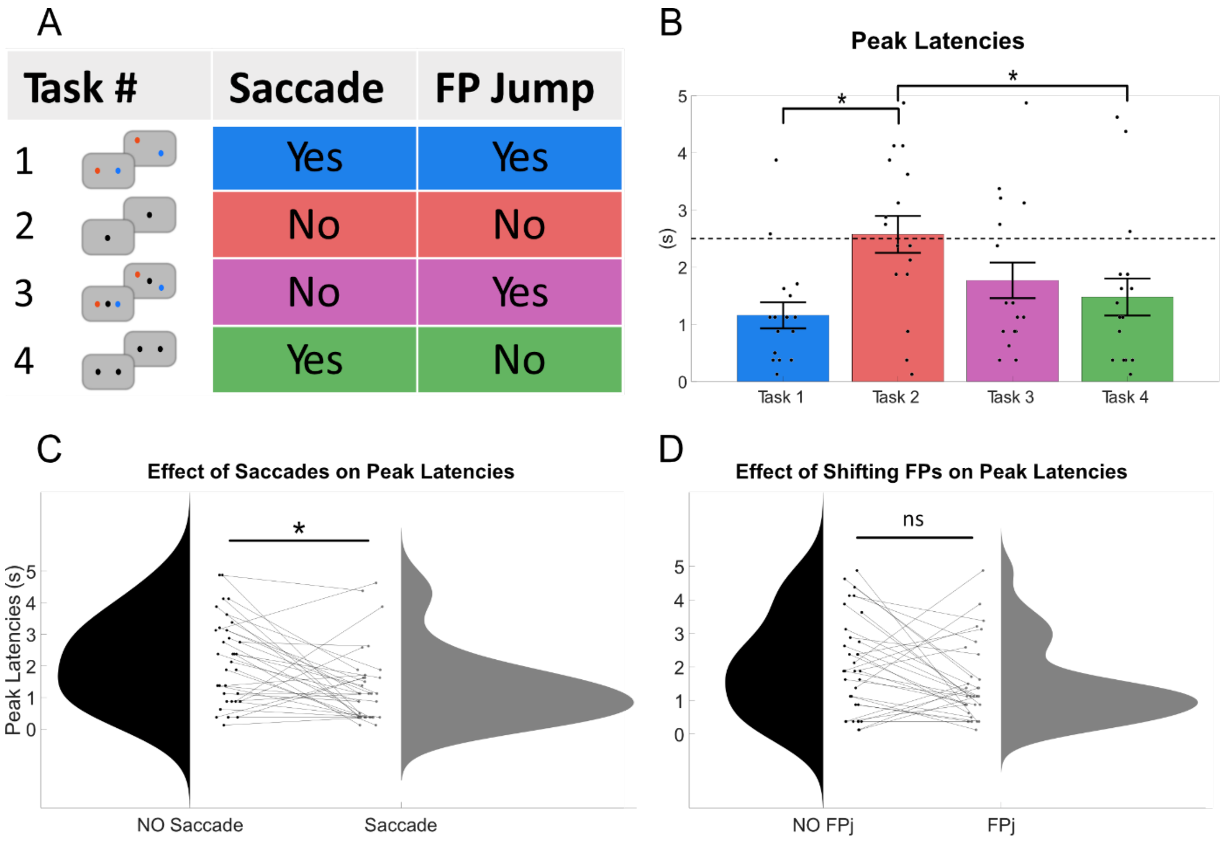


**Supplementary Figure 8: Across-task comparison of peak PSLs**. Each latency data point on **(B-C-D)** is obtained from the PSL distribution of an individual as the latency bin with the highest probability value. **(B)** Across task peak PSL comparisons reveal that task identity has a significant effect on the PSL bin with the highest probability. Task pairs that significantly differed from each other after correction for multiple comparisons are shown with asterisks. When grouped according to the presence of saccades **(C)** and jumping fixation points **(D)**, we found a main effect of saccades on the peak PSLs, and no effect of shifts in fixation point locations.

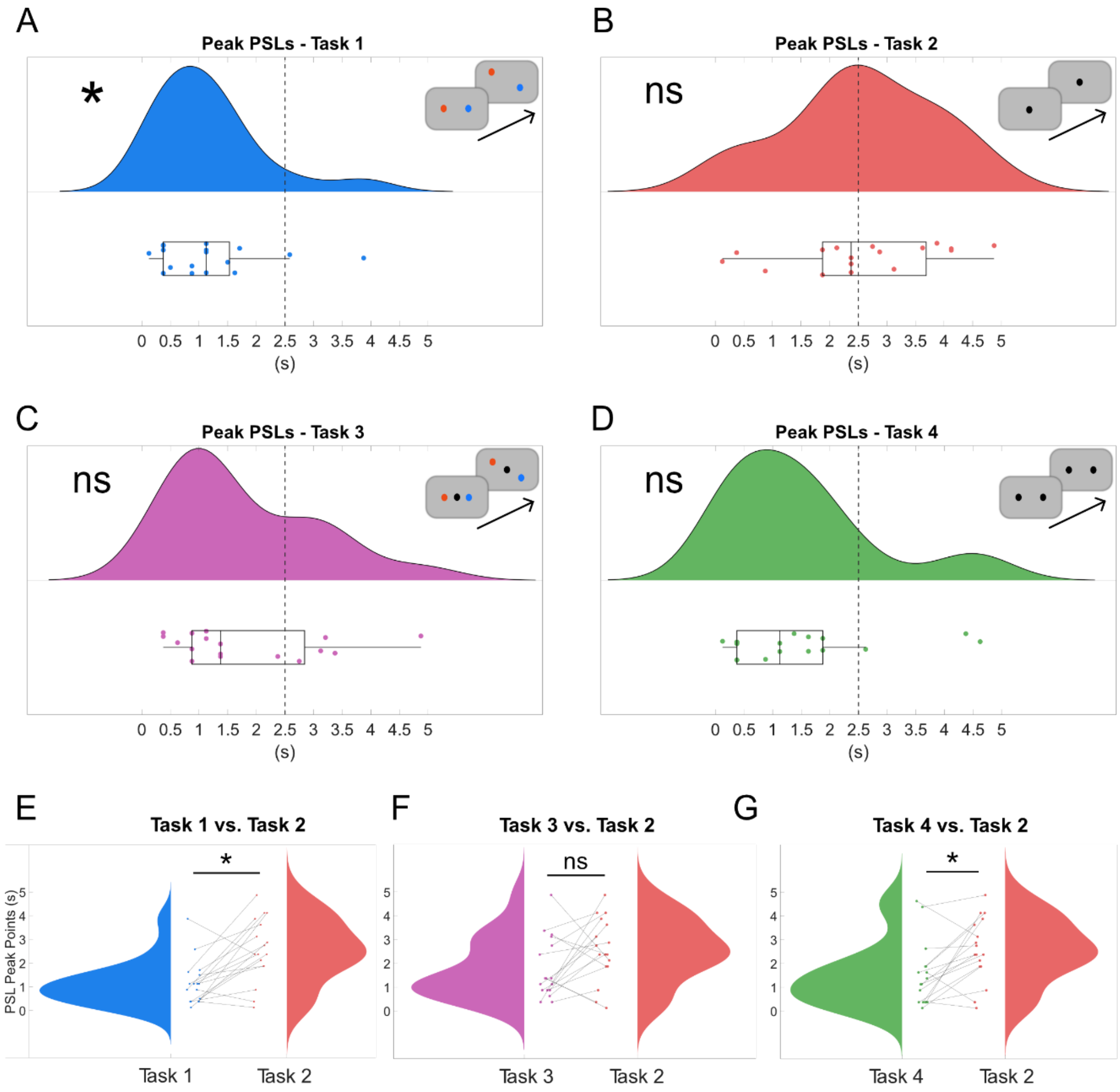


**Supplementary Figure 9: Single-task (A-D) and task-pair (E-G) comparisons of peak PSLs**. Wilcoxon signed-rank tests against the mid-point of trials showed that peak PSLs are significantly closer to trial onsets in Task 1, but not in Tasks 2,3, and 4, despite showing a leftward trend of peaks in Tasks 3 and 4. **(E-G)** In comparison with Task 2, subjects have significantly smaller peak PSL values in Tasks 1 and 4, but not in Task 3.

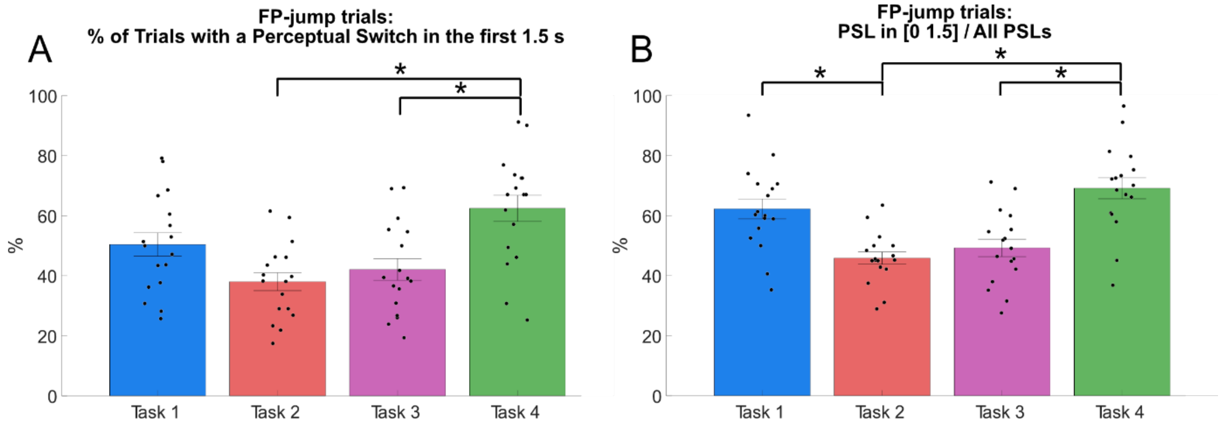


**Supplementary Figure 10**: **Analysis of the first 1.5 seconds following trial onsets with a shift in fixation point locations. (A)** Across-task comparison of % trials that included a perceptual switch within the first 1.5 seconds, only using trials where the fixation points shifted at trial onset. Task identity has an overall main effect on this fraction. Task 4 has a significantly higher percentage of trials with a perceptual switch in the first 1.5 seconds than Task 2 and Task 3. **(B)** Across-task comparison of % of PSLs below 1.5 seconds, only using trials where the fixation points shifted at trial onset. Task identity was found to affect this fraction significantly, and the post-hoc tests showed that the fraction differs significantly between Tasks 1 vs. 2, 2 vs. 4, and 3 vs. 4 (shown with asterisks).

### Supplementary Tables

#### Supplementary Table 1: Basic information regarding both Physical (PHYS) and binocular rivalry (BR) trials.

| | # of PHYS Trials | | | | % Success in PHYS Trials | | | | # of BR Trials | | | | # of Completed Switches (BR) | | | | # of Incomplete Switches (BR) | | | | Average Perceptual Dominance Durations in BR (s) | | | |
|---|---|---|---|---|---|---|---|---|---|---|---|---|---|---|---|---|---|---|---|---|---|---|---|---|
| **Subj** | Task 1 | Task 2 | Task 3 | Task 4 | Task 1 | Task 2 | Task 3 | Task 4 | Task 1 | Task 2 | Task 3 | Task 4 | Task 1 | Task 2 | Task 3 | Task 4 | Task 1 | Task 2 | Task 3 | Task 4 | Task 1 | Task 2 | Task 3 | Task 4 |
| 1 | 104 | 104 | 104 | 104 | 100.00 | 97.12 | 97.12 | 97.12 | 91 | 91 | 91 | 91 | 61 | 57 | 60 | 86 | 12 | 14 | 11 | 7 | 5.63 | 5.38 | 5.71 | 4.85 |
| 2 | 104 | 104 | 104 | 104 | 100.00 | 100.00 | 97.12 | 99.04 | 91 | 91 | 91 | 91 | 45 | 25 | 27 | 54 | 47 | 40 | 27 | 30 | 4.28 | 5.19 | 5.62 | 5.18 |
| 3 | 104 | 40* | 104 | 104 | 100.00 | NaN | 100.00 | 97.12 | 91 | 35* | 91 | 91 | 83 | NaN | 82 | 116 | 16 | NaN | 12 | 1 | 4.15 | NaN | 4.61 | 4.00 |
| 4 | 104 | 104 | 104 | 104 | 100.00 | 99.04 | 100.00 | 100.00 | 91 | 91 | 91 | 91 | 115 | 121 | 123 | 142 | 4 | 1 | 2 | 20 | 3.34 | 3.35 | 3.42 | 2.72 |
| 6 | 104 | 104 | 104 | 104 | 100.00 | 92.31 | 100.00 | 100.00 | 91 | 91 | 91 | 91 | 118 | 98 | 120 | 160 | 10 | 15 | 32 | 57 | 3.16 | 3.66 | 3.00 | 2.33 |
| 8 | 104 | 104 | 104 | 104 | 100.00 | 99.04 | 100.00 | 95.19 | 91 | 91 | 91 | 91 | 89 | 80 | 72 | 88 | 9 | 0 | 1 | 4 | 4.11 | 4.98 | 5.26 | 4.84 |
| 9 | 104 | 104 | 104 | 104 | 100.00 | 97.12 | 91.35 | 100.00 | 91 | 91 | 91 | 91 | 93 | 86 | 65 | 142 | 33 | 25 | 33 | 41 | 3.47 | 3.87 | 4.24 | 2.62 |
| 10 | 96 | 104 | 104 | 104 | 100.00 | 100.00 | 100.00 | 100.00 | 84 | 91 | 91 | 91 | 48 | 74 | 62 | 79 | 37 | 30 | 19 | 57 | 4.57 | 4.10 | 5.26 | 3.65 |
| 11 | 112 | 96 | 104 | 96 | 79.46 | 98.96 | 95.19 | 95.83 | 98 | 84 | 91 | 84 | 62 | 92 | 86 | 106 | 13 | 4 | 9 | 1 | 5.02 | 4.05 | 4.45 | 3.93 |
| 12 | 104 | 104 | 104 | 104 | 100.00 | 100.00 | 100.00 | 100.00 | 91 | 91 | 91 | 91 | 58 | 72 | 46 | 115 | 10 | 1 | 3 | 3 | 5.51 | 5.28 | 7.39 | 3.73 |
| 13 | 104 | 104 | 104 | 104 | 100.00 | 98.08 | 98.08 | 100.00 | 91 | 91 | 91 | 91 | 69 | 95 | 68 | 137 | 10 | 1 | 3 | 7 | 4.90 | 3.91 | 5.61 | 3.18 |
| 14 | 104 | 104 | 104 | 104 | 99.04 | 100.00 | 100.00 | 100.00 | 91 | 91 | 91 | 91 | 90 | 46 | 81 | 159 | 37 | 22 | 30 | 45 | 3.57 | 6.16 | 3.64 | 2.45 |
| 15 | 104 | 104 | 104 | 104 | 97.12 | 100.00 | 100.00 | 99.04 | 91 | 91 | 91 | 91 | 120 | 116 | 129 | 174 | 27 | 15 | 19 | 49 | 2.79 | 3.11 | 2.99 | 2.22 |
| 16 | 104 | 104 | 104 | 104 | 100.00 | 100.00 | 99.04 | 100.00 | 91 | 91 | 91 | 91 | 75 | 69 | 69 | 145 | 0 | 1 | 0 | 3 | 4.44 | 5.13 | 5.40 | 2.85 |
| 17 | 104 | 104 | 104 | 104 | 100.00 | 100.00 | 100.00 | 100.00 | 91 | 91 | 91 | 91 | 49 | 31 | 40 | 85 | 38 | 34 | 35 | 37 | 4.53 | 5.65 | 5.30 | 3.51 |
| 18 | 104 | 104 | 104 | 104 | 100.00 | 100.00 | 100.00 | 97.12 | 91 | 91 | 91 | 91 | 138 | 103 | 122 | 162 | 5 | 16 | 24 | 5 | 2.73 | 3.28 | 2.60 | 2.61 |
| 19 | 104 | 104 | 104 | 104 | 100.00 | 100.00 | 100.00 | 100.00 | 91 | 91 | 91 | 91 | 62 | 53 | 60 | 75 | 7 | 2 | 12 | 2 | 5.43 | 7.06 | 5.63 | 5.32 |
| 20 | 104 | 104 | 104 | 104 | 100.00 | 98.08 | 100.00 | 100.00 | 91 | 91 | 91 | 91 | 35 | 59 | 65 | 61 | 5 | 4 | 1 | 0 | 6.65 | 5.23 | 5.98 | 5.64 |
| **Mean (SD)** | **104.00 (2.74)** | **103.50 (1.94)** | **104.00 (0.00)** | **103.60 (1.89)** | **98.65 (4.84)** | **98.81 (1.97)** | **98.77 (2.33)** | **98.91 (1.65)** | **91.00 (2.40)** | **90.59 (1.70)** | **91.00 (0.00)** | **90.61 (1.65)** | **78.33 (29.52)** | **75.12 (27.76)** | **76.50 (29.61)** | **115.90 (38.01)** | **17.78 (14.52)** | **13.24 (13.06)** | **15.17 (12.46)** | **20.50 (21.52)** | **4.35 (1.06)** | **4.67 (1.11)** | **4.78 (1.26)** | **3.65 (1.12)** |

Asterisks indicates that Subject #3 was excluded from further analysis due to significantly lower trial counts than any other subject in Task 2.

**Supplementary Table 2: Median PSLs of each individual, noted as [Median (IQR)].**

| | PSL median (IQR) - PHYS | | | | PSL median (IQR) - BR complete | | | | PSL median (IQR) - BR incomplete | | | |
|---|---|---|---|---|---|---|---|---|---|---|---|---|
| Subj no. | Task 1 | Task 2 | Task 3 | Task 4 | Task 1 | Task 2 | Task 3 | Task 4 | Task 1 | Task 2 | Task 3 | Task 4 |
| 1 | 0.64 (0.52) | 0.50 (0.30) | 0.80 (0.86) | 1.41 (1.10) | 2.76 (2.45) | 2.50 (2.54) | 2.17 (3.08) | 1.85 (1.93) | 2.86 (2.98) | 2.57 (2.80) | 3.43 (0.97) | 1.69 (2.16) |
| 2 | 0.51 (0.35) | 0.51 (0.09) | 0.48 (0.16) | 0.59 (0.25) | 1.31 (1.77) | 2.86 (2.77) | 3.38 (2.59) | 1.72 (2.41) | 1.78 (1.62) | 1.87 (1.71) | 2.03 (2.79) | 1.06 (0.72) |
| 3 | 0.80 (0.80) | NaN (NaN) | 0.63 (0.35) | 1.19 (0.97) | 2.32 (3.17) | NaN (NaN) | 2.50 (2.53) | 2.16 (2.02) | 2.09 (1.61) | NaN (NaN) | 2.35 (2.30) | 0.98 (0.00) |
| 4 | 0.78 (0.81) | 0.56 (2.10) | 0.57 (0.55) | 1.24 (0.51) | 2.15 (2.85) | 2.45 (2.37) | 2.27 (2.41) | 2.62 (2.76) | 1.66 (2.83) | 0.24 (0.00) | 3.30 (0.15) | 1.31 (0.39) |
| 6 | 0.62 (0.67) | 0.47 (0.31) | 0.60 (0.52) | 0.57 (1.26) | 2.54 (2.69) | 2.62 (2.38) | 2.19 (2.50) | 2.52 (2.62) | 1.03 (0.82) | 2.47 (2.23) | 2.98 (1.95) | 2.27 (3.16) |
| 8 | 0.74 (0.47) | 0.59 (0.27) | 0.56 (0.18) | 1.01 (1.22) | 2.19 (2.65) | 2.73 (2.42) | 2.29 (2.30) | 1.78 (2.51) | 1.56 (2.60) | NaN (NaN) | 3.68 (0.00) | 1.51 (1.02) |
| 9 | 0.54 (0.27) | 0.40 (0.18) | 0.38 (0.16) | 0.49 (0.45) | 3.18 (1.94) | 2.46 (2.13) | 1.95 (2.70) | 2.14 (2.50) | 3.29 (3.23) | 2.36 (2.12) | 2.22 (2.22) | 1.32 (2.64) |
| 10 | 0.59 (0.30) | 0.54 (0.14) | 0.56 (0.15) | 0.54 (0.16) | 1.72 (3.26) | 2.39 (1.98) | 1.53 (2.45) | 2.73 (2.94) | 2.07 (2.77) | 2.65 (3.33) | 2.64 (1.59) | 0.62 (0.81) |
| 11 | 1.28 (2.26) | 0.52 (1.35) | 0.93 (1.65) | 0.85 (0.64) | 2.29 (2.28) | 2.51 (2.25) | 2.87 (2.26) | 1.44 (2.44) | 1.48 (2.29) | 3.29 (1.15) | 2.83 (1.95) | 0.88 (0.00) |
| 12 | 0.67 (0.68) | 0.59 (0.55) | 0.58 (0.13) | 0.78 (0.97) | 1.50 (1.68) | 2.27 (2.23) | 1.46 (1.67) | 1.47 (2.58) | 0.76 (0.41) | 3.92 (0.00) | 1.93 (1.02) | 0.57 (0.22) |
| 13 | 0.84 (1.45) | 0.52 (1.38) | 0.42 (0.11) | 0.68 (1.00) | 2.07 (2.66) | 2.79 (2.33) | 1.79 (3.01) | 2.42 (2.54) | 2.87 (2.47) | 2.08 (0.00) | 0.41 (0.23) | 1.52 (1.26) |
| 14 | 0.52 (0.35) | 0.41 (0.12) | 0.44 (1.03) | 0.44 (0.28) | 2.47 (3.07) | 2.00 (3.04) | 2.67 (2.32) | 2.69 (2.96) | 1.34 (3.00) | 1.79 (2.07) | 1.10 (2.82) | 1.07 (3.07) |
| 15 | 0.54 (0.63) | 0.54 (0.66) | 0.56 (2.30) | 0.55 (0.79) | 3.13 (3.26) | 2.46 (2.54) | 2.30 (2.44) | 2.87 (3.55) | 0.46 (0.58) | 3.30 (2.22) | 2.91 (1.58) | 0.39 (1.24) |
| 16 | 0.64 (0.23) | 0.46 (0.08) | 0.47 (0.53) | 1.17 (0.64) | 2.18 (2.41) | 2.68 (2.48) | 2.49 (2.63) | 2.00 (2.20) | NaN (NaN) | 0.98 (0.00) | NaN (NaN) | 1.38 (0.94) |
| 17 | 0.55 (0.20) | 0.54 (0.24) | 0.64 (0.99) | 1.05 (0.24) | 0.56 (1.77) | 2.46 (1.45) | 1.73 (2.29) | 0.93 (1.74) | 0.52 (0.92) | 2.40 (2.80) | 2.28 (2.82) | 0.93 (0.56) |
| 18 | 0.88 (1.64) | 0.57 (0.91) | 0.59 (1.78) | 1.39 (0.75) | 2.86 (2.86) | 2.57 (2.26) | 2.26 (2.62) | 2.49 (2.41) | 3.43 (1.85) | 2.18 (2.86) | 1.74 (1.68) | 3.29 (1.71) |
| 19 | 0.78 (0.46) | 0.60 (0.15) | 0.67 (0.24) | 1.35 (0.53) | 2.10 (2.33) | 2.94 (2.40) | 2.51 (1.92) | 2.08 (1.57) | 3.02 (2.45) | 4.23 (1.14) | 3.04 (2.18) | 1.87 (1.71) |
| 20 | 0.76 (0.20) | 0.53 (0.26) | 0.49 (0.32) | 1.29 (0.30) | 1.31 (1.47) | 2.36 (2.24) | 2.90 (2.59) | 2.01 (1.62) | 1.13 (1.29) | 1.38 (2.09) | 1.31 (0.00) | NaN (NaN) |
| **Mean (SD)** | **0.71 (0.19)** | **0.52 (0.06)** | **0.58 (0.14)** | **0.92 (0.35)** | **2.15 (0.68)** | **2.53 (0.23)** | **2.29 (0.49)** | **2.11 (0.52)** | **1.84 (0.96)** | **2.36 (1.03)** | **2.36 (0.88)** | **1.33 (0.70)** |

**Supplementary Table 3: Trials with FP-jump, detected trial onset-locked saccades in those trials, and rate of saccades that putatively "cause" a perceptual switch in 10 subjects with reliable saccade detection.**

| Subj | # Trials with FP jump | | # Detected Saccades | | | # Saccades w/Perceptual Switch in the same trial | | | # Saccades Followed by a perceptual switch within 0.7 sec | | # Saccades Followed by a perceptual switch within 1 sec | |
|---|---|---|---|---|---|---|---|---|---|---|---|---|
| | Task 1 | Task 4 | Task 1 | Task 2 | Task 4 | Task 1 | Task 2 | Task 4 | Task 1 | Task 2 | Task 4 | Task 2 |
| 2 | 68 | 91 | 22 | 91 | 50 | 11 | 20 | 33 | 6 | 4 | 19 | 5 |
| 8 | 71 | 91 | 29 | 91 | 31 | 23 | 53 | 22 | 9 | 9 | 11 | 13 |
| 9 | 69 | 91 | 16 | 91 | 28 | 13 | 60 | 11 | 3 | 11 | 5 | 19 |
| 10 | 70 | 91 | 11 | 91 | 39 | 6 | 50 | 17 | 6 | 8 | 15 | 11 |
| 12 | 72 | 91 | 25 | 91 | 72 | 11 | 50 | 47 | 6 | 13 | 35 | 18 |
| 13 | 76 | 91 | 18 | 91 | 45 | 5 | 56 | 32 | 1 | 18 | 21 | 21 |
| 14 | 60 | 91 | 16 | 91 | 51 | 10 | 32 | 25 | 5 | 6 | 20 | 9 |
| 15 | 73 | 91 | 24 | 91 | 21 | 12 | 63 | 5 | 7 | 14 | 5 | 22 |
| 16 | 70 | 91 | 38 | 91 | 68 | 27 | 47 | 47 | 12 | 7 | 36 | 8 |
| 20 | 78 | 91 | 28 | 91 | 68 | 8 | 41 | 35 | 3 | 12 | 20 | 14 |
| **Median** | **70.5** | **91** | **23** | **91** | **47.5** | **11** | **50** | **28.5** | **6** | **10** | **19.5** | **13.5** |
| **Total** | **707** | **910** | **227** | **910** | **473** | **126** | **472** | **274** | **58** | **102** | **187** | **140** |
| **% of all detected saccades** | | | | | | **55.5** | **51.9** | **57.9** | **25.5** | **11.2** | **39.5** | **15.4** |
